# Chloroplast genome-based genetic resources for Japan’s threatened subalpine forests via genome skimming

**DOI:** 10.1101/2023.12.03.569577

**Authors:** James R.P. Worth, Satoshi Kikuchi, Seiichi Kanetani, Daiki Takahashi, Mineaki Aizawa, Elena A. Marchuk, Hyeok Jae Choi, Maria A. Polezhaeva, Viktor V. Sheiko, Saneyoshi Ueno

## Abstract

The Japanese subalpine zone is dominated by a distinct and ecologically important conifer rich forest biome, subalpine coniferous forests, that are an outlier of the extensive boreal forests of Eurasia. While being relatively intact compared to other forest biomes in Japan, subalpine coniferous forests are under significant threat from deer browsing, global warming and small populations size effects. However, there is a severe lack of genetic resources available for the study of this biome’s major constituent plant species. This study aims to develop chloroplast genome-based genetic resources for 12 widespread subalpine tree and shrub species via genome skimming of whole genomic DNA using short reads (100-150 bp in length). For 10 species, whole chloroplast genomes were assembled via *de novo*-based methods from 4-10 individuals per species sampled from across their ranges in Japan and, for non-Japanese endemic species, elsewhere in northeast Asia. A total of 566 single nucleotide polymorphisms for Japanese samples and 768 for all samples (varying from 2 to 202 per species) were identified which were distributed in geographically restricted lineages in most species. In addition, between 9 to 58 polymorphic simple sequence repeat regions were identified per species. For two Ericaceae species (*Rhododendron brachycarpum* and *Vaccinium vitis-idaea*) characterized by large chloroplast genomes, *de novo* assembly failed, but single nucleotide polymorphisms could be identified using reference mapping. This data will be useful for genetic studies of the taxonomic relationship of populations within Japan and to other parts of northeast Asia, investigating phylogeographic patterns within species, conservation genetics and has potential application for studies of environmental and ancient DNA.

## Introduction

Genome skimming is the shallow sequencing of genomic DNA enabling the accurate sequencing of genomic components such as the chloroplast, mitochondria and high copy nuclear regions. In the last decade genome skimming has revolutionised the ease with which genomic data can be obtained for both model and non-model species (Dodsworth, 2015). For example, the ability to cost effectively and reliably assemble the relatively small mitochondria of animals and chloroplast genome of plants using *de novo* or reference mapping methods from low coverage genome skimming data has accelerated our understanding of the variation in size, gene content and arrangement of these genomes (Li *et al*., 2020) with thousands of whole organelle genomes now available on Genbank. Whole mitochondria (De Mandal *et al*., 2014) and chloroplast (Song *et al*., 2023) genomes are now being incorporated into barcode libraries (called ultra barcodes) providing greater power to understand the phylogenetic affinity of taxa and increase the ability to identify the small fragments of organelle DNA derived from ancient or environmental samples. The relative low cost of genome skimming means that the whole organelle genome of multiple samples can feasibly be obtained, facilitating the discovery of potentially phylogenetically informative characters and mutational hotspots for the taxonomic level of interest (e.g. genus or species level). In plants, such studies have increased in recent years identifying informative chloroplast DNA variation for phylogenetic (Malé *et al*., 2014; Fu *et al*., 2022), phylogeographic (Migliore, Lézine and Hardy, 2020) and conservation genetic (Worth *et al*., 2023) studies.

Here we use genome skimming of genomic DNA to develop chloroplast genome genetic resources for an important but threatened forest biome in Japan, subalpine coniferous forests. These forests, classified in the Abieti-Piceetalia jesoensis order (Krestov and Omelko, 2010), are an outlier of the extensive boreal forests of Eurasia. In Japan, they have a wide but fragmented distribution from northern Hokkaido (45.4° N) to the highest peaks of the Kii Peninsula (34.1° N) and Shikoku Island (33.7° N) in western Japan (Figure 1). The fossil record shows that subalpine forests were more widespread during glacial periods, occurring more than a thousand metres below their current elevational limit to near sea level in Honshu, but contracted their ranges during warm interglacials involving significant range losses, especially in western Japan (e.g. the loss of *Tsuga diversifolia* (Maxim.) Mast.) from Chugoku (Takahara, Tanida and Miyoshi, 2001), *Pinus koraiensis* Siebold et Zucc from most of western Japan (Aizawa, Kim and Yoshimaru, 2012) and *Picea maximowiczii* Regel ex Mast. from Kyushu and Chugoku (Magri *et al*., 2020)). Japan’s subalpine coniferous forests are of immense value for biodiversity, harbouring many endemic species, and for the ecosystem services they provide, for example, via their role as the headwater forests of Japan’s major watersheds. Although significant areas of subalpine forests have been replaced by Japanese larch (*Larix kaempferi* (Lamb.) Carrière) plantations (Franklin *et al*., 1979), the biome is one of the comparatively least disturbed by humans. However, despite this these forests face significant threats including severe browsing by increasing populations of deer, small population size effects and rising global temperatures (Oishi and Doei, 2015; Tsuyama *et al*., 2015). Subalpine coniferous forests are considered most at risk of decline in western Japan where they have little available higher ground to migrate to already being restricted to near the tops (Hämet-Ahti, Ahti and Koponen, 1974) of relatively low mountain ranges. Indeed, populations of many species of subalpine trees and shrubs in western Japan are endangered (Japanese Red Data Book online: http://jpnrdb.com/) especially in Shikoku and the Kii Peninsula.

**Figure 1.**
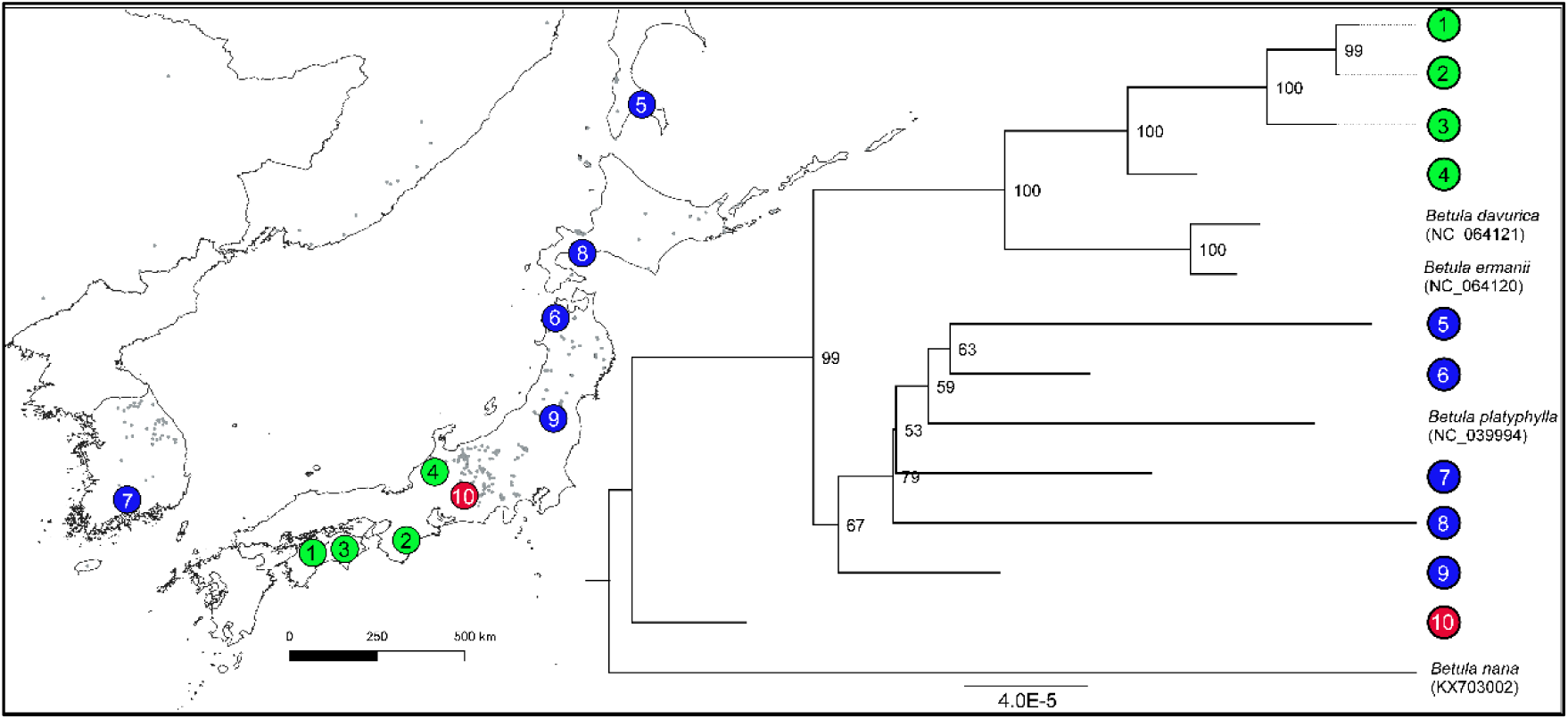
Phylogenetic tree based on the whole chloroplast genomes of *Betula ermanii* assembled from samples from Japan, South Korea and Russia. The only other *Betula ermanii* whole chloroplast genome sequence available on Genbank and three outgroup genomes are also included. The location of the identified clades are mapped.

Genetic markers can inform conservation management of plants by resolving taxonomic uncertainty of species, identify populations which should be prioritised for conservation based on their genetic distinctiveness (Shibabayashi *et al*., 2023) and reveal areas with significantly high or low genetic diversity. However, there are limited genetic markers available for the component species of Japan’s subalpine forests. The existing range-wide organelle-based genetic studies (*Abies veitchii* (Uchiyama *et al*., 2023), *Picea jezoensis* (Aizawa *et al*., 2007), *Pinus koraiensis* (Aizawa, Kim and Yoshimaru, 2012) and *Vaccinium vitis-idea* (Ikeda *et al*., 2015)) all are based on Sanger sequencing of a few chloroplast fragments. In this study we develop genetic resources based on whole chloroplast genome sequencing via genome skimming for 12 trees and shrubs (Table 1) that are key components of Japan’s subalpine forests occurring either as canopy dominants or important in the understory (Yamanaka, 1959; Franklin *et al*., 1979; Sugita, 1992) and have ranges extending from northern to western Japan. The 12 species include five conifers (*Abies veitchii* Lind., *Picea jezoensis* (Siebold et Zucc.) Carrière, *Pinus koraiensis*. *Thuja standishii* (Gordon) Carrière and *Tsuga diversifolia*) and seven angiosperms (*Acer ukurunduense* Trautv. et C.A.Mey., *Berberis amurensis* Rupr., *Betula ermanii* Cham., *Ilex rugosa* F.Schmidt, *Oplopanax japonicus* Nakai, *Rhododendron brachycarpum* D.Don ex G.Don and *Vaccinium vitis-idaea* L.). Four (*A. veitchii*, *O. japonicus*, *T. diversifolia* and *T. standishii*) are endemic to Japan while the remaining eight species occur in other parts of Eurasia including Korea, northeast China and Far East Russia (Song, 1991; Krestov and Nakamura, 2002). For 8 of 12 species the whole chloroplast genome is available but these consist of single samples collected from their ranges outside Japan (e.g. in China, *B. ermanii*, *A. ukurunduense*, and South Korea *B. amurensis*, *V. vitis-idaea*, *P. koraiensis*, *P. jezoensis* or in Japan, *T. diversifolia* and *T. standishii*). No whole chloroplast genome is available for *O. japonicus*, *R. brachycarpum*, *I. rugusa* or *A. veitchii*. Here we report intraspecific chloroplast genome diversity (single nucleotide polymorphisms (SNPs) and indels including simple sequence repeats (SSRs)) based on whole chloroplast genome sequencing of these species for the first time, identify how this intraspecific diversity is geographically distributed and, lastly, assess how the diversity identified in this study is related to already published genomes of the same species.

**Table 1.** Details of the 12 subalpine plants investigated in this study including their taxonomy, distribution and habit.

| Species | Family | Distribution | Habit |
| --- | --- | --- | --- |
| *Abies veitchii* | Pinaceae | Japanese endemic | Tree |
| *Acer ukurunduense* | Sapindaceae | Japan/ Korea/ Far E. Russia/ N.E. China | Tree |
| *Berberis amurensis* | Berberidaceae | Japan/ Korea/ Far E. Russia/ N.E. China | Shrub |
| *Betula ermanii* | Betulaceae | Japan/ Korea/ Far E. Russia/ N.E. China | Tree |
| *Ilex rugosa* | Aquifoliaceae | Japan/ Far E. Russia | Shrub |
| *Oplopanax japonicus* | Araliaceae | Japanese endemic | Shrub |
| *Picea jezoensis* | Pinaceae | Japan/ Korea/ Far E. Russia/ N.E. China | Tree |
| *Pinus koraiensis* | Pinaceae | Japan/ Korea/ Far E. Russia/ N.E. China | Tree |
| *Rhododendron brachycarpum* | Ericaceae | Japan/ Korea/ Far E. Russia | Shrub |
| *Thuja standishii* | Cupressaceae | Japanese endemic | Tree |
| *Tsuga diversifolia* | Pinaceae | Japanese endemic | Tree |
| *Vaccinium vitis-idaea* | Ericaceae | Northern hemisphere | Shrub |

## Materials and Methods

### Sampling and DNA extraction

Leaf samples from four to twelve individuals of each of the twelve study species were collected from across their ranges in Japan and where possible where they occur outside Japan in N.E. China, South Korea and Far East Russia (for a list of all samples see Table S1 in Supplementary Materials). In Japan the sampling included areas where the species are widespread in central Honshu but also the most northern populations in Hokkaido/ Tohoku and most southern parts of the species ranges in western Japan. DNA was extracted using either the DNeasy Plant Mini Kit (Qiagen) or a modified CTAB protocol. DNA concentration and quality were assessed by agarose gel electrophoresis and a Qubit 2.0 fluorometer (Life Technologies).

### Chloroplast genome assembly

Whole genomic DNA was sent to the Beijing Genomic Institute where short-size libraries were constructed and paired-end sequencing (2 x 150 bp except for *T. diversifolia* which were 2 x 100 bp) was performed on an Illumina HiSeq2000 Genome Analyser. Chloroplast genomes were assembled using GetOrganelle v1.7.7.0 (Jin *et al*., 2020) and its dependencies including SPAdes (Bankevich *et al*., 2012) and Bowtie2 (Langmead and Salzberg, 2012), using default setting, or, in the case of *Abies veitchii*, Novoplasty 4.3.1 (Dierckxsens, Mardulyn and Smits, 2016) was used as it was found to perform more reliably for this species. The previously published whole chloroplast genome of *Tsuga diversifolia* (Genbank accession: MH171102) was re-assembled using GetOrganelle. The chloroplast genomes were annotated using GeSeq (Tillich *et al*., 2017) with annotations checked by aligning to reference genomes and prepared for submission to Genbank using GB2sequin (Lehwark and Greiner, 2019).

The chloroplast genomes of *R. brachycarpum* and *V. vitis-idaea* (both Ericaceae) could not be assembled using GetOrganelle or Novoplasty despite trying a range of settings. This was likely caused by an unsolvable tangled graph due to long repeats (Fu *et al*., 2022) with Ericaceae chloroplast genomes being characterized by gene rearrangement, repetitive regions and IR expansion (Li *et al*., 2020) making them difficult to assemble using short Illumina reads alone. Instead reads were mapped to reference chloroplast genomes with ‘bwa mem’ (Li, 2013). *Vaccinium vitis-idaea* reads were mapped to a *V. vitis-idaea* whole chloroplast genome (sourced from a plant cultivated in Canada) assembled from Oxford Nanopore (SRR25468450) and Illumina NovaSeq reads (SRR25477290) (Hirabayashi, Debnath and Owens, 2023) using the ptGAUL pipeline (Zhou *et al*., 2023) (https://github.com/Bean061/ptgaul). Before chloroplast genome assembly, Nanopore reads were filtered by Filtlong (https://github.com/rrwick/Filtlong) with parameters of ‘--min_length 1000 --keep_percent 90,’ while Illumina reads (1 Gbp for each pair) were down-sampled by the ‘sample’ subcommand of seqtk (https://github.com/lh3/seqtk) ver. 1.4-r122 with the parameter of ‘-s 0’. A chloroplast genome of the same species from Korea (Genbank accession: LC521968) was not used because it was found to be missing approximately 5,000 bp of sequence. For *R. brachycarpum,* no reference of the same species, or raw data including long nanopore reads suitable to assemble a reference genome, were available. Therefore, the reads were mapped to the closest available whole chloroplast genome on Genbank as determined by blasting the longest chloroplast contigs produced by GetOrganelle. This resulted in the chloroplast genome of *R. calophytum* Franch. (OM373082.1 (Ma *et al*., 2022)) and *R. shanii* W.P. Fang (MW374796 (Yu *et al*., 2022)) being the closest. bcftools ver. 1.17 (Li, 2011) was used to create consensus sequences of each mapping incorporating both SNPs and indel sites whereby the mapping consensus had the reference sequence in regions where SNPs/indels were not called for all samples or the minimum read depth of one of the samples was < 20 or above an average depth + one standard deviation. The read mapping-based assemblies were not annotated or submitted to Genbank due to the relatively high number of ambiguous sites compared to the *de novo* method and the incorporation of reference sequence in regions with minimum/maximum read depths over the designated thresholds. Rather they were used primarily for uncovering SNPs and discovery of the main lineages in these species.

### Phylogenetic analyses

For the ten species where whole chloroplast *de novo* assembly was successful, the intraspecific phylogenetic relationships of the chloroplast genomes were investigated using maximum likelihood implemented in RAxML v.8.2.10 (Stamatakis, 2014). The input files were prepared in Geneious 9.15 (Biomatters, New Zealand) where the whole chloroplast genomes were aligned using MAFFT multiple aligner Version: 1.3.6 (Katoh *et al*., 2002). For use as outgroups, complete published chloroplast genomes of the same species (if available) and up to three complete genomes of different species within the same genus as determined by Blastn (Altschul *et al*., 1990) (i.e. the three with the closest percentage identity) were used. This was done due to the fact that in some cases the chloroplast genomes of the most closely related species were unavailable and, for the angiosperms, the fact that chloroplast relationships do not necessarily follow species relationships (e.g. for *Betula* (Palme *et al*., 2004), *Acer* (Saeki *et al*., 2011)). Before alignment one inverted repeat region was removed for all angiosperms and poorly aligned regions excluded using Gblocks 0.91b (Castresana, 2000) ran with default setting except with gaps allowed. The same method as above was used for phylogenetic tree construction for species assembled by reference mapping, *R. brachycarpum* and *V. vitis-idaea*, except that no outgroups of congeneric species were used.

The accuracy of the whole chloroplast genome sequencing was checked using Sanger sequencing of variations identified in six species (3 angiosperms and 3 conifers) five of whose chloroplast genome was obtained via GetOrganelle *de novo* method and *R. brachycarpum* obtained using reference mapping. A total of 25 primer pairs in six species were designed around variable sites identified in the genome skimming based whole chloroplast genome data using the Primer3 2.3.4. plugin (Untergasser *et al*., 2012) in Geneious and Sanger sequenced using Supredye (MS Techno Systems) following the manufacturer’s instructions for two samples per species. Sequences were determined via capillary electrophoresis on an ABI3130 Genetic Analyzer (Life Technologies, Waltham, MA, USA).

### Genetic diversity

The whole chloroplast genomes of each species were aligned in Geneious using MAFFT (Katoh *et al*., 2002) including accessions of the same species available on Genbank where available (except *Pinus koraiensis* (NC_004677) and *Thuja standishii* (KX832627) where the records on Genbank were found to have unusually high genetic distance from other whole chloroplast genomes of the same species obtained both from Genbank and/or this study and were therefore excluded).

The number of non-informative sites (singletons), parsimony informative sites (i.e. sites that are present at least twice and are therefore potentially informative for phylogenetic analyses), overall nucleotide diversity and average number of nucleotide differences (*k*) were calculated in DNAsp v.6.12.03 (Rozas *et al*., 2017). The number of indel events and indel diversity was calculated using the multi-allelic gap option in the same program. Calculations were done separately for those samples from Japan and for all samples (i.e. individuals sampled for this study within and outside Japan and obtained from Genbank). In addition, the number of SNPs and nucleotide diversity of genes and intergenic spacers across the whole chloroplast genomes of each species was calculated in DNAsp v.6.12.03 using the multi-domain function.

Due to the presence of ambiguous sites in the reference mapping-based chloroplast genome assemblies of *R. brachycarpum* and *V. vitis-idaea* that is not compatible with DNAsp v.6.12.03 we used PopART (Leigh and Bryant, 2015) to calculate the number of SNPs, parsimony informative sites and overall nucleotide diversity.

### Chloroplast microsatellite identification

For the ten species where whole chloroplast *de novo* assembly was successful chloroplast microsatellite regions were searched for using Find Polymorphic SSRs which uses the Phobos Tandem Repeat Finder (Mayer, 2008) in Geneious with a repeat unit length of 1-3 bp, a minimum length of 10 bp and a requirement that they occur in all sequences in the alignment.

## Results

### Chloroplast genome assembly

An average of 8,285,799 reads were obtained per sample for the 12 species (Table S2). For the 10 species that had their whole chloroplast genome successfully *de novo* assembled, the percentage of total reads that were from the chloroplast genome varied between 0.24% to 6.64% (Table S2) with substantial difference between angiosperms, average of 3.46%, and the large nuclear genome bearing conifers with an average of 0.90%. As a consequence, overall read coverage of the chloroplast genome was 266.0 for angiosperms to 92.8 for conifers (Table S2). Whole chloroplast genome lengths were between 156,109 to 166,708 bp for angiosperms and between 116,924 to 130,925 bp for conifers. The whole chloroplast genomes of each species did not differ substantially in length (average difference = 185.8 bp) with the most similar being *I. rugosa* (6 bp maximum difference between samples) and the most different being *B. amurensis* (431 bp maximum difference). All sequences obtained from direct Sanger sequencing were identical to the chloroplast genomes (except for some ambiguous sites in the read mapping consensuses of *R. brachycarpum*) and confirmed all the variable sites. For the list of Genbank accession numbers for each sample see Table S2 and for *R. brachycarpum* and *V. vitis-idaea* sequence data see the fasta alignments in the Supplementary Materials.

### *Genetic diversity of* de novo *assembled species*

An average of 56.6 SNPs per species were discovered when considering only Japanese samples (Table 2) and 76.8 when including all samples (Table 2 and S3). A total of 31% of all SNPs were parsimony informative for Japan and 41% for all samples. For both angiosperms and conifers, the number of SNPs for samples from Japan varied greatly between species with highest values 138 for *B. amurensis* and 110 for *P. jezoensis* and lowest being 2 for *T. standishii* and 25 for *I. rugosa*, respectively (Table 2). When considering all samples, the number of SNPs increased by between 6.4 to 1.07 times (average 2.21 times) with the highest increase in *A. ukurunduense* and *P. jezoensis* (Table S3). An average of 61.5 indels events were observed in Japanese samples with a maximum of 147 in *B. ermanii* and a low of 17 in *T. standishii*. Similar to SNPs, the number of indel events increased when including all samples (average 83.8 per species). Considering SSR type indels for Japanese samples, mono-repeats were more common than either di- or tri- repeats in all species with an average of 42.8 per species ranging from 72 in *B. amurensis* to 22 in *T. standishii* (Table 3). This compares to an average per species of 9.3 for di- and 6.4 for tri- repeats. An average of 54.9% of mono-repeats were polymorphic varying from 90.2% in *A. veitchii* to 0% in *T. standishii*. This contrasts to an average of 7.5% and 4.7% per species being polymorphic for di- and tri- repeats, respectively. When considering all samples, the number of overall mono-, di- and tri-repeats increased slightly with the number of polymorphic ones increasing to 59.7%, 8.06% and 7.4% percent for mono-, di- and tri- repeats, respectively (Table S4).

**Table 2.** Summary of genetic diversity identified in the 12 study species including single nucleotide polymorphisms and indels based on the Japan dataset. Total sites analysed is the length of the whole chloroplast genome after excluding one inverted repeat.

| Species | No. samples | Total sites | Sites (excluding gaps) | Monomorphic sites | Singletons | Parsimony informative sites | Nucleotide diversity ($P_i$) | Average number of nucleotide differences ($k$) | Indel events | InDel Diversity per site ($P_i(i)$) |
| --- | --- | --- | --- | --- | --- | --- | --- | --- | --- | --- |
| *Acer ukurunduense* | 6 | 130025 | 129681 | 129671 | 3 | 7 | 0.00004 | 4.7 | 31 | 0.00011 |
| *Berberis amurensis* | 8 | 129781 | 128946 | 128808 | 95 | 43 | 0.00036 | 46.1 | 139 | 0.00041 |
| *Betula ermanii* | 8 | 134983 | 134079 | 134008 | 47 | 24 | 0.00018 | 24.4 | 147 | 0.00039 |
| *Ilex rugosa* | 9 | 131506 | 131476 | 131451 | 17 | 8 | 0.00006 | 7.6 | 26 | 0.00008 |
| *Oplopanax japonicus* | 8 | 130338 | 130185 | 130125 | 6 | 54 | 0.00022 | 29 | 29 | 0.0001 |
| *Abies veitchii* | 7 | 121579 | 121109 | 121046 | 57 | 6 | 0.00016 | 19.8 | 100 | 0.00031 |
| *Picea jezoensis* | 4 | 124559 | 124047 | 123937 | 103 | 7 | 0.00046 | 56.5 | 53 | 0.00022 |
| *Pinus koraiensis* | 4 | 117154 | 116760 | 116731 | 26 | 3 | 0.00013 | 15 | 37 | 0.00017 |
| *Thuja standishii* | 5 | 131166 | 130225 | 130223 | 2 | 0 | 0.000006 | 0.8 | 17 | 0.00006 |
| *Tsuga diversifolia* | 4 | 121291 | 120827 | 120769 | 34 | 24 | 0.00027 | 33 | 36 | 0.00017 |
| *Rhododendron brachycarpum*<sup>1</sup> | 7 | 157211 | - | 157122 | 47 | 42 | 0.00023 | - | - | - |
| *Rhododendron brachycarpum*<sup>2</sup> | 7 | 155954 | - | 155875 | 46 | 33 | 0.00019 | - | - | - |
| *Vaccinium vitis-idaea*<sup>3</sup> | 10 | 144322 | - | 144109 | 129 | 84 | 0.00043 | - | - | - |

<sup>1</sup> mapped to MW374796; <sup>2</sup> mapped to OM373082; <sup>3</sup> mapped to the *V. vitis-idaea* whole chloroplast genome assembled from the Oxford Nanopore and Illumina NovaSeq reads Hirabayashi, Debnath and Owens (2023)

**Table 3.** Summary of genetic diversity identified at simple sequence repeat (SSR) regions identified in the *de novo* assembled whole chloroplast genomes of 10 study species for Japan samples only.

| Species | No.<br>samples | No.<br>polymorphic<br>SSRs<br>(mono-repeats >9<br>length) | No.<br>monomorphic<br>SSRs (mono-repeats >9<br>length) | No.<br>polymorphic<br>SSRs (di-<br>repeats >9<br>length) | No.<br>monomorphic<br>SSRs (di-<br>repeats >9<br>length) | No.<br>polymorphic<br>SSRs (tri-repeats >9<br>length) | No.<br>monomorphic<br>SSRs (tri-repeats >9<br>length) |
| --- | --- | --- | --- | --- | --- | --- | --- |
| *Acer ukurunduense* | 6 | 21 | 47 | 0 | 3 | 0 | 12 |
| *Berberis amurensis* | 8 | 53 | 19 | 0 | 9 | 2 | 2 |
| *Betula ermanii* | 8 | 44 | 17 | 2 | 16 | 1 | 11 |
| *Ilex rugosa* | 9 | 21 | 30 | 0 | 4 | 0 | 5 |
| *Oplopanax japonicus* | 8 | 9 | 18 | 0 | 7 | 0 | 2 |
| *Abies veitchii* | 7 | 37 | 4 | 1 | 13 | 0 | 9 |
| *Picea jezoensis* | 4 | 17 | 8 | 4 | 7 | 0 | 5 |
| *Pinus koraiensis* | 4 | 20 | 16 | 0 | 6 | 0 | 0 |
| *Thuja standishii* | 5 | 0 | 22 | 0 | 14 | 0 | 9 |
| *Tsuga diversifolia* | 4 | 13 | 12 | 0 | 7 | 0 | 6 |

Based on the Japan only data, an average of 85.2% of regions (genes and intragenic spacers) in the angiosperm chloroplast and 86.8% of regions in the conifer chloroplast were invariable with most of the variable regions only having one SNP (Figure S1 and S2). The results for all samples are not shown because they are nearly identical to those based on the only Japan data. For the majority of species, the most diverse regions based on nucleotide diversity were intragenic spacers except for the low diversity *A. ukurunduense* in Japan where half of the most diverse regions were within genes (Table 4 and 5). The most diverse regions based on nucleotide diversity were particular to each species in 56 cases, 13 regions were found in at least two species and only three regions, trnH-GUG̶̶─psbA, ycf1─ndhF and the long ycf1 gene, were observed in three species and none in four or more. For the results based on all samples see Table S5.

**Table 4.**
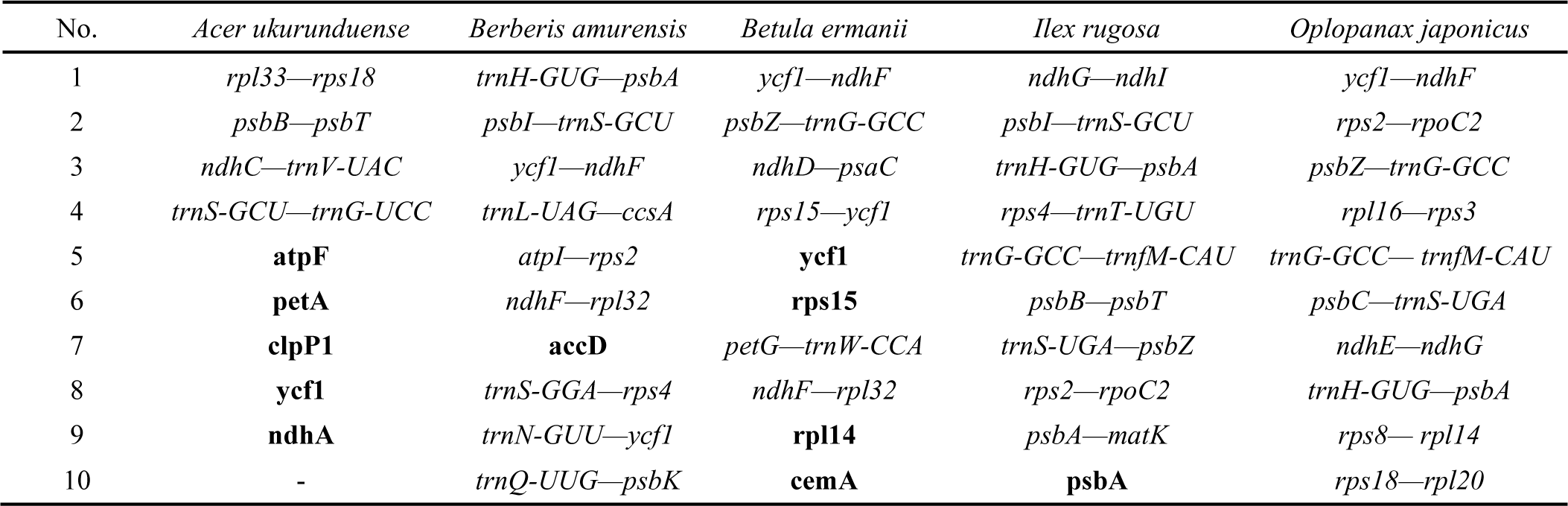
The top ten most variable regions of the *de novo* assembled whole chloroplast genome of Japanese samples of five angiosperm study species according to nucleotide diversity (*Pi*). Intergenic spacers are italicized while genes are shown in bold. Note that *Acer ukurunduense* only had nine variable regions.

**Table 5.** The top ten most variable regions of the *de novo* assembled whole chloroplast genome of Japanese samples of five conifer study species according to nucleotide diversity (*Pi*). Intergenic spacers are italicized while genes are shown in bold. Note that *Thuja standishii* only had two variable regions.

| No. | *Abies veitchii* | *Picea jezoensis* | *Pinus koraiensis* | *Thuja standishii* | *Tsuga diversifolia* |
| --- | --- | --- | --- | --- | --- |
| 1 | *ycf2—trnH-GUG* | *rpoA—rps11* | *rps12—rps7* | *ndhJ—rps12* | *clpP—ycf12* |
| 2 | *rpl33—psaJ* | **trnR—UCU** | *trnfM-CAU—trnG-GCC* | *chlB—matK* | *rps12—rps7* |
| 3 | *atpE—trnM-CAU* | **trnS—UGA** | **psbK** | - | *rps11—rpl36* |
| 4 | *trnI-CAU—psbA* | *rpoC1—rpoC2* | *rrn5—trnR-ACG* | - | *psbJ—psbL* |
| 5 | *psaJ—trnP-UGG* | *chlB—trnQ-UUG* | *trnS-GCU—psaM* | - | *ycf2—trnH-GUG* |
| 6 | *trnP-UGG—trnW-CCA* | *ycf12—clpP* | *trnQ-UUG—psbK* | - | *chlN—ycf1* |
| 7 | *clpP—rps12* | *trnH-GUG—trnT-GGU* | *rpl23—trnI-GAU* | - | **rps15** |
| 8 | **psbK** | *psaJ—trnP-UGG* | *trnT-GGU—trnS-GCU* | - | **rpl14** |
| 9 | *trnM-CAU—trnV-UAC* | *ycf12—psbB* | *psbK—psbI* | - | *psbK—psbI* |
| 10 | *rpl23—trnV-GAC* | *ycf1—rps15* | *chlN—rps15* | - | **ycf1** |

### Genetic diversity of reference mapping assembled species

The number of overall SNPs identified in *V. vitis-idaea* for was 213 for Japanese samples with 84 being parsimony informative. For all samples the number of SNPs increased to 238 with 84 parsimony informative. In *R. brachycarpum*, the mapping using the *R. shanii* reference chloroplast genome (MW374796) resulted in a higher number of SNPs being uncovered (89/ 107 overall SNPs with 42/ 54 parsimony informative for Japan and all samples, respectively) compared to mapping to *R. calophytum* (OM373082) which uncovered 79 and 98 overall SNPs with 33 and 44 parsimony informative for only Japan and all samples, respectively.

### Phylogenetic relationships

The chloroplast variation in most species was found to be distributed in well supported clades with clear non-overlapping geographical ranges (see Figures 1-12). For example, *B. ermanii*, *A. ukurunduense*, *O. japonicus*, *P. jezoensis*, *R. brachycarpum*, *V. vitis-idaea* and (to a less clear extent) *B. amurensis* were found to harbour northern and southern distributed lineages. These lineages coincided with the distribution of two varieties in the case of *P. jezoensis*, var. *jezoensis* distributed Far East Russia, northeast China and Hokkaido and var. *hondoensis* distributed in Honshu, Japan (Aizawa *et al*., 2007). For *R. brachycarpum* phylogenetic relationships were similar between the results based on the two references used for read mapping with all main clades recovered in both but the phylogeny based on the *R. calophytum* reference was less well resolved in the early diverging branches probably due to a lower number of SNPs recovered (Figure 11 and S3). On the other hand, the lineages identified in *P. koraiensis* and *I. rugosa* had overlapping ranges while for *A. veitchii* a diverged lineage was observed in only one individual at the southern edge of the species range (Figure 1). For both *T. diversifolia* and *T. standishii* all individuals harboured genetically similar chloroplast genomes with no diverged lineages (Figure 1). Genbank accessions of the same species were mostly placed within one of the distinct clades identified in this study (e.g. *P. koraensis*, A. *ukurunduense*, and *B. amurensis*). In three species, *A. veitchii*, *B. ermanii* and *B. amurensis*, Genbank accessions of outgroup species were nested within identified clades.

**Figure 2.**
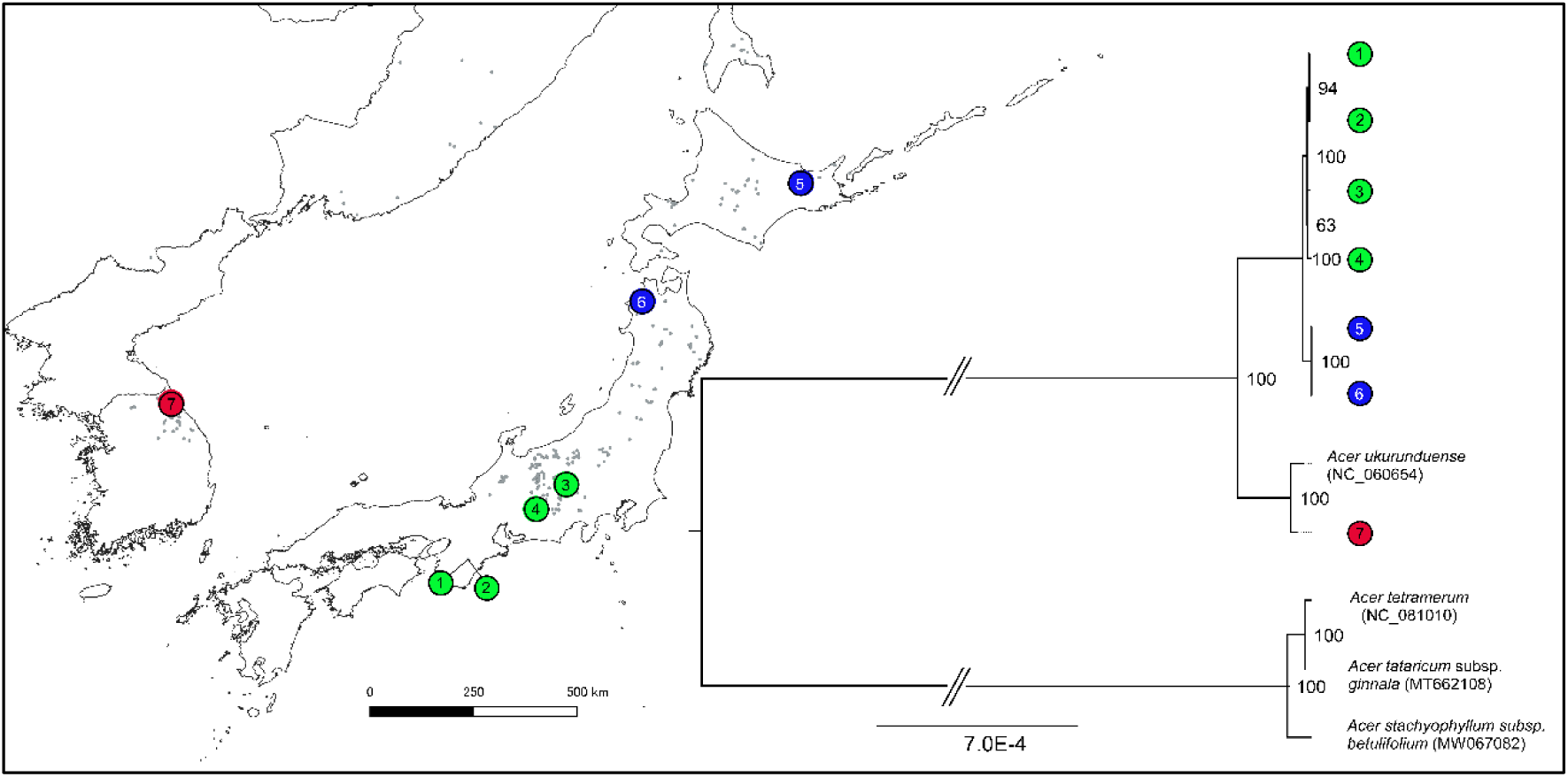
Phylogenetic tree based on the whole chloroplast genomes of *Acer ukurunduense* assembled from samples from Japan and South Korea. The only other *Acer ukurunduense* whole chloroplast genome sequence available on Genbank and three outgroup genomes are also included. The location of the identified clades are mapped.

**Figure 3.**
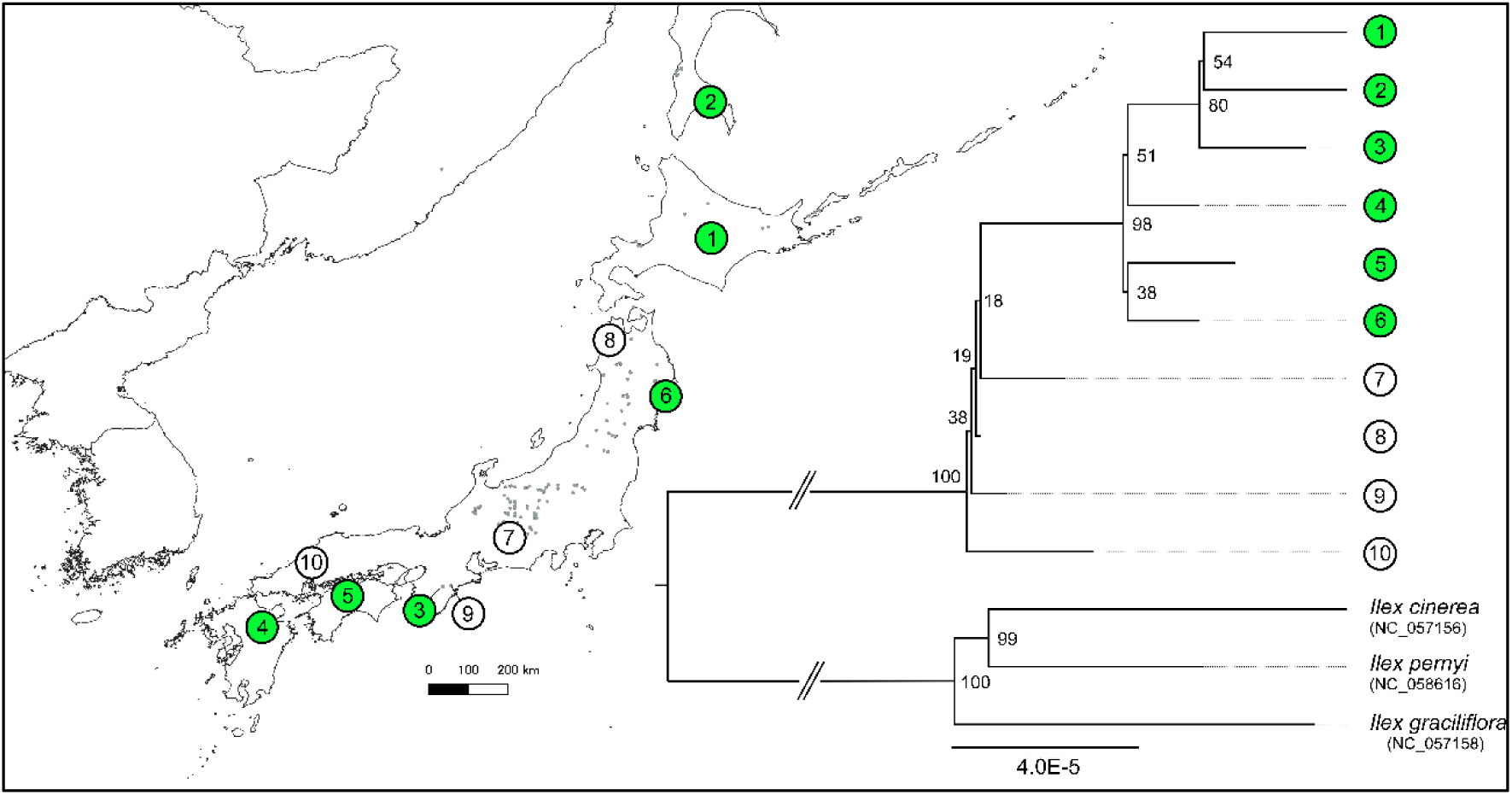
Phylogenetic tree based on the whole chloroplast genomes of *Ilex rugosa* assembled from samples from Japan and Russia. Three outgroup genomes available on Genbank are also included. The location of the identified clades are mapped.

**Figure 4.**
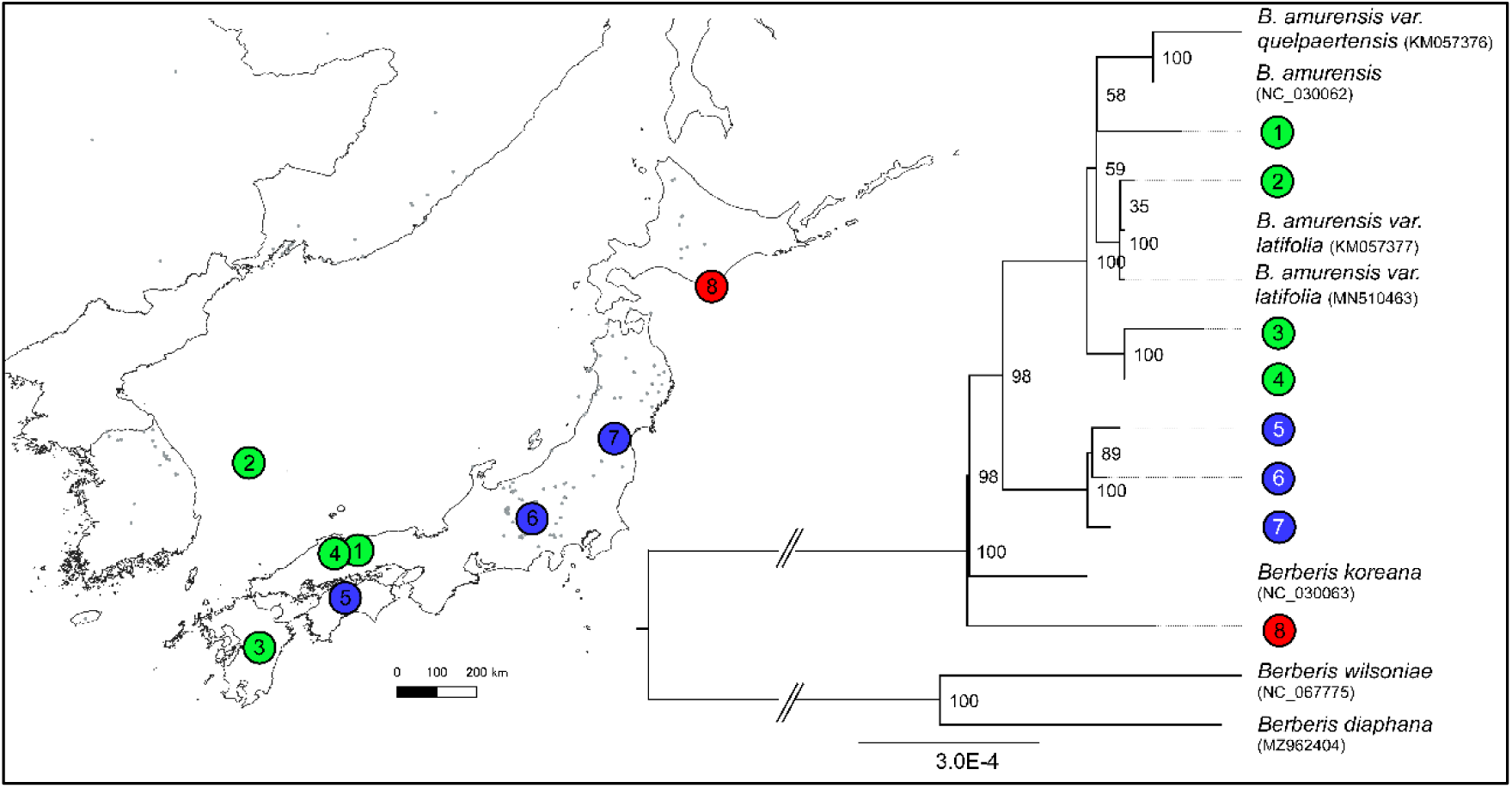
Phylogenetic tree based on the whole chloroplast genomes of *Berberis amurensis* assembled from samples from Japan and South Korea. Other *Berberis amurensis* whole chloroplast genome sequences available on Genbank and three outgroup genomes are also included. The location of the identified clades are mapped.

**Figure 5.**
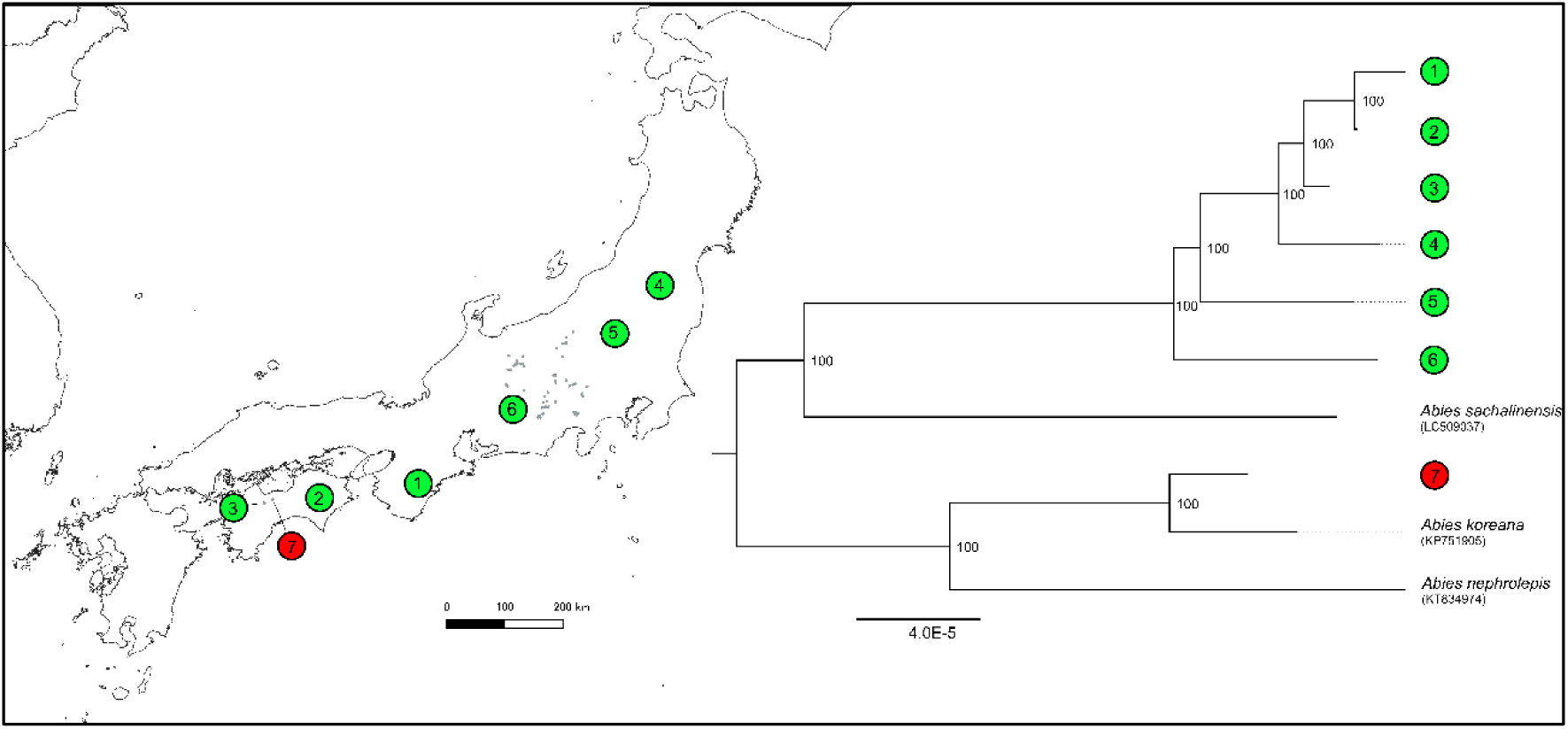
Phylogenetic tree based on the whole chloroplast genomes of *Abies veitchii* assembled from samples from Japan. Three outgroup genomes available on Genbank are also included. The location of the identified clades are mapped.

**Figure 6.**
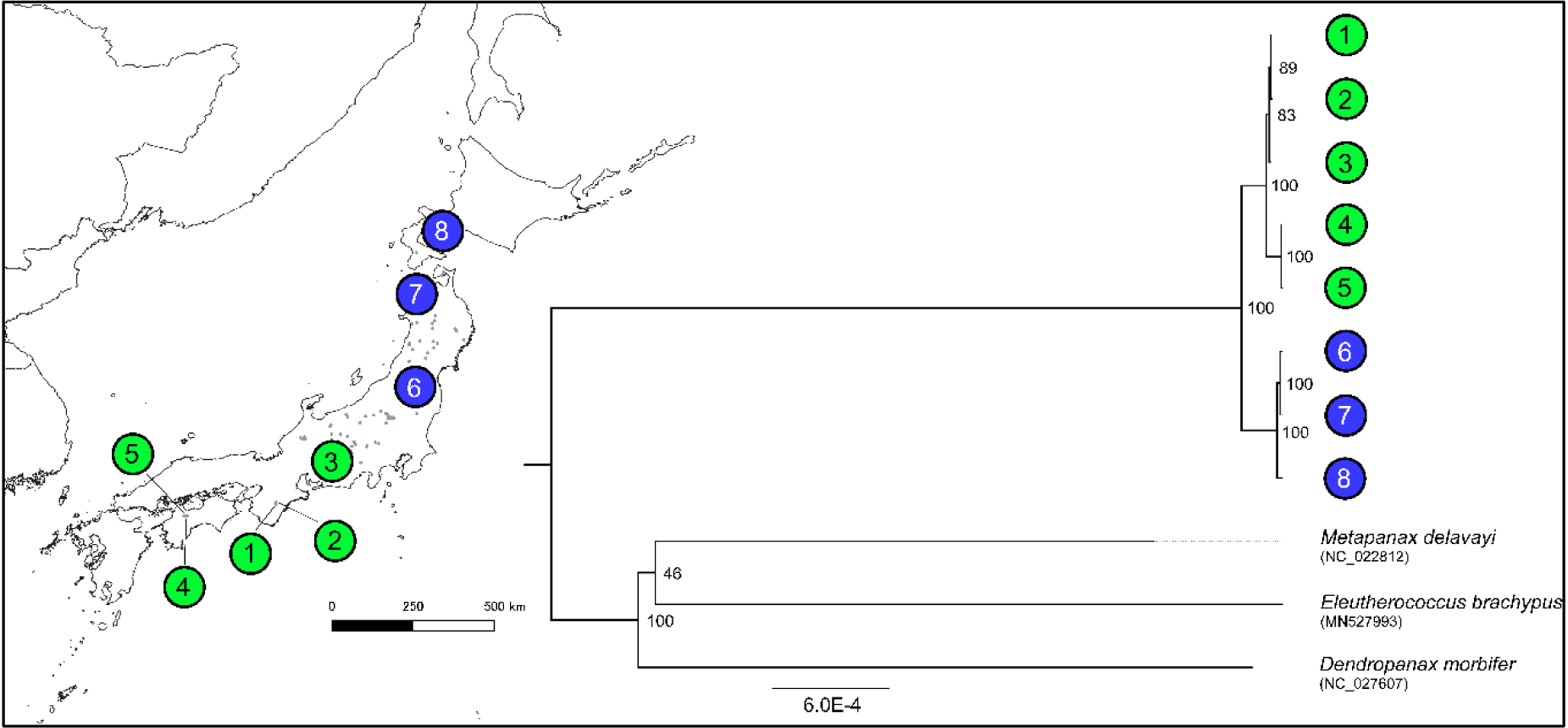
Phylogenetic tree based on the whole chloroplast genomes of *Oplopanax japonicus* assembled from samples from Japan. Three outgroup genomes available on Genbank are also included. The location of the identified clades are mapped.

**Figure 7.**
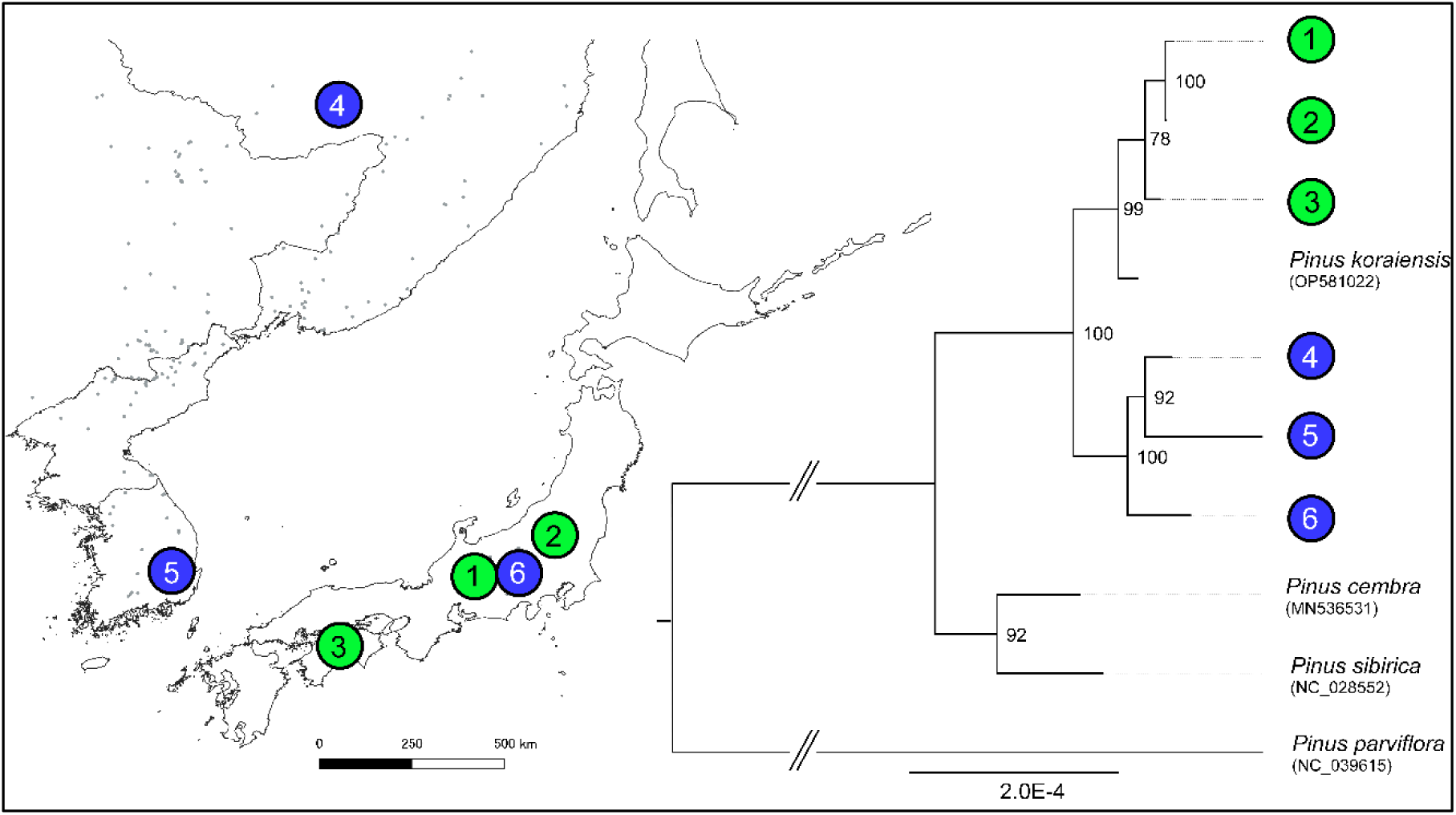
Phylogenetic tree based on the whole chloroplast genomes of *Pinus koraiensis* assembled from samples from Japan, South Korea and Russia. One other *Pinus koraiensis* whole chloroplast genome sequence available on Genbank and three outgroup genomes are also included. The location of the identified clades are mapped.

**Figure 8.**
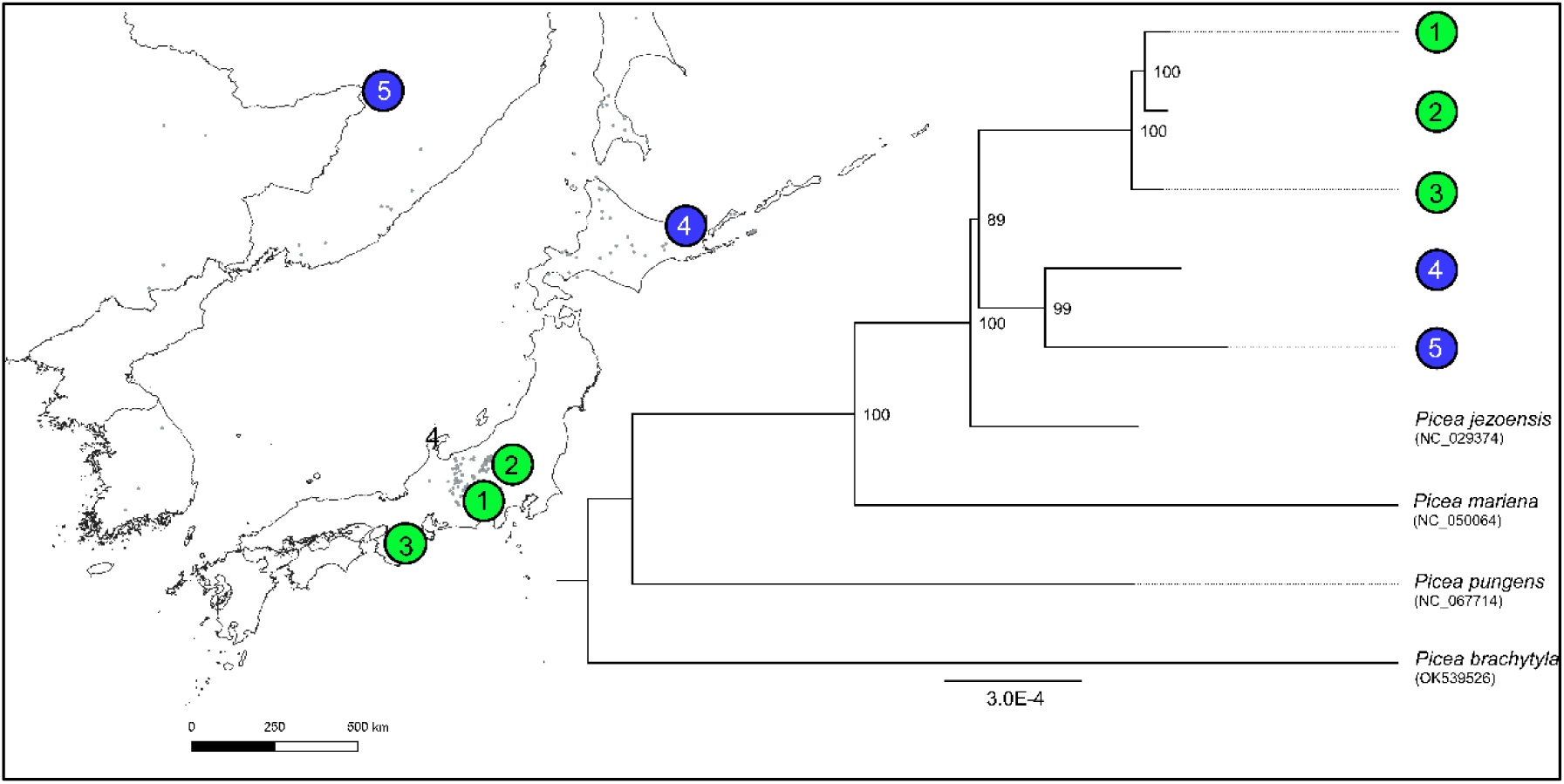
Phylogenetic tree based on the whole chloroplast genomes of *Picea jezoensis* assembled from samples from Japan and Russia. One other *Picea jezoensis* whole chloroplast genome sequence available on Genbank and three outgroup genomes are also included. The location of the identified clades are mapped.

**Figure 9.**
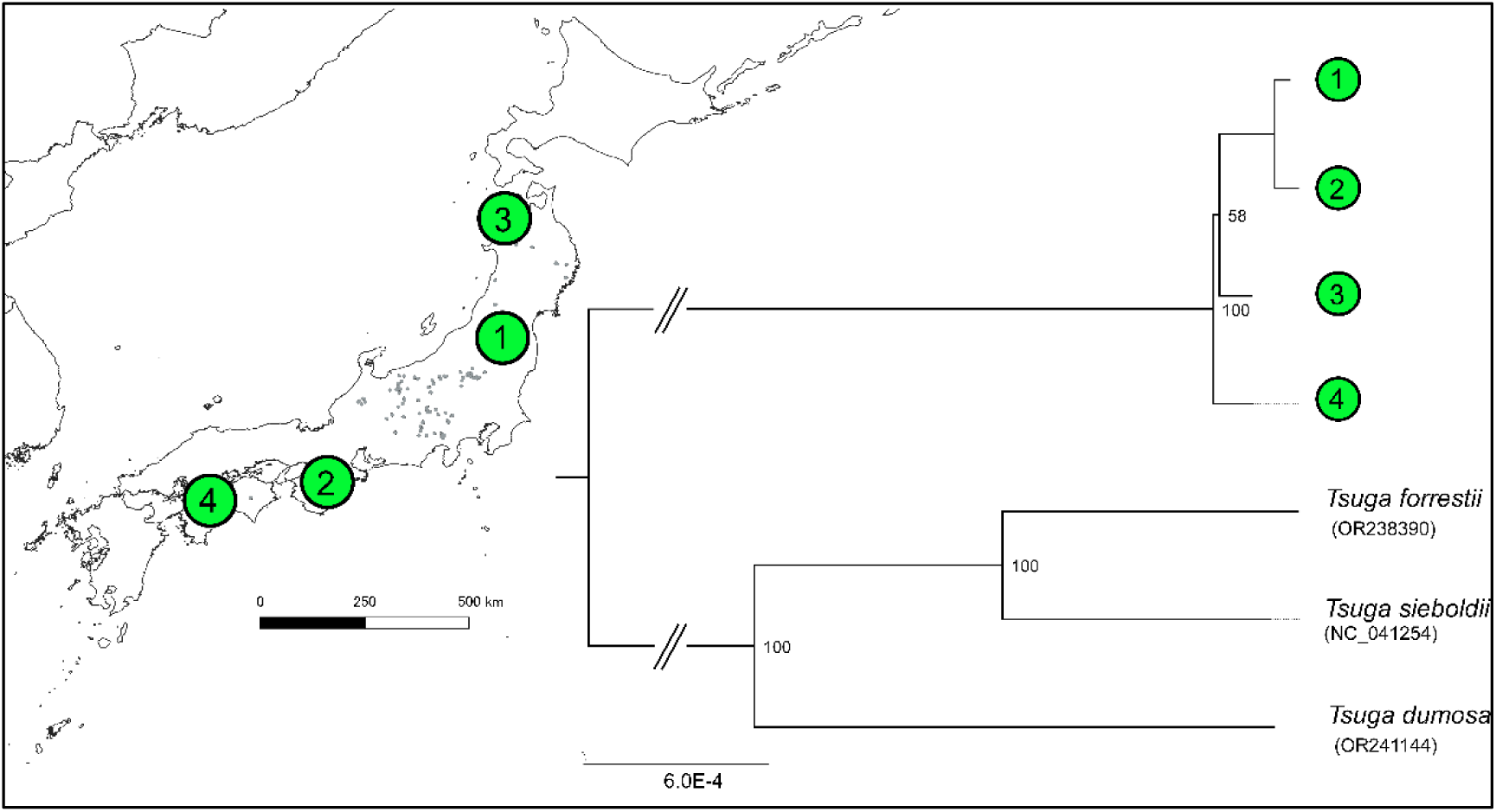
Phylogenetic tree based on the whole chloroplast genomes of *Tsuga diversifolia* assembled from samples from Japan. Three outgroup genomes available on Genbank are also included. The location of the identified clades are mapped.

**Figure 10.**
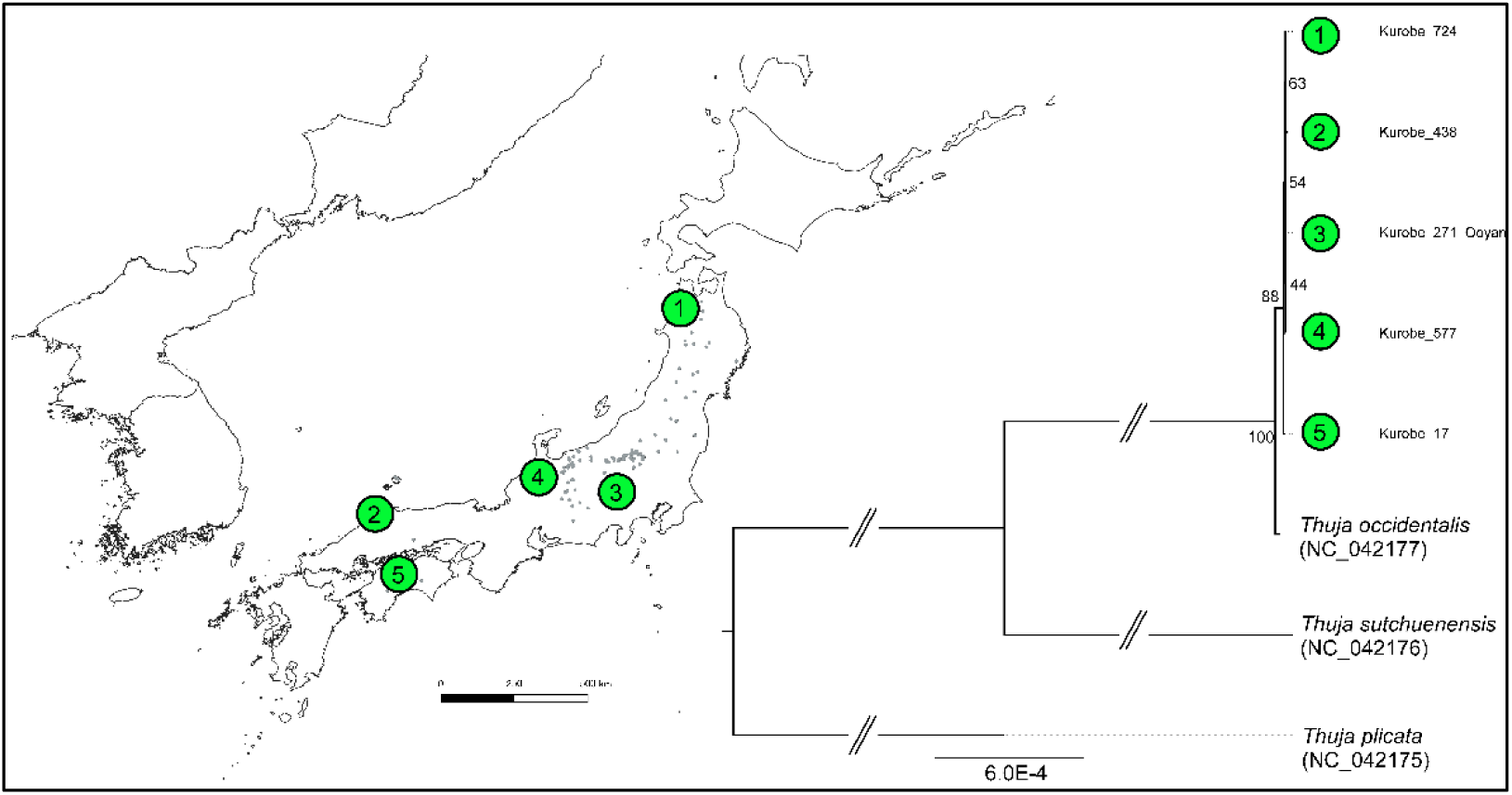
Phylogenetic tree based on the whole chloroplast genomes of *Thuja standishii* assembled from samples from Japan. Three outgroup genomes available on Genbank are also included. The location of the identified clades are mapped. Note that the outgroup sequences may not reliably represent the true phylogenetic relationships of *T. standishii* to other *Thuja* species but here are only used for the purpose of providing outgroups.

**Figure 11.**
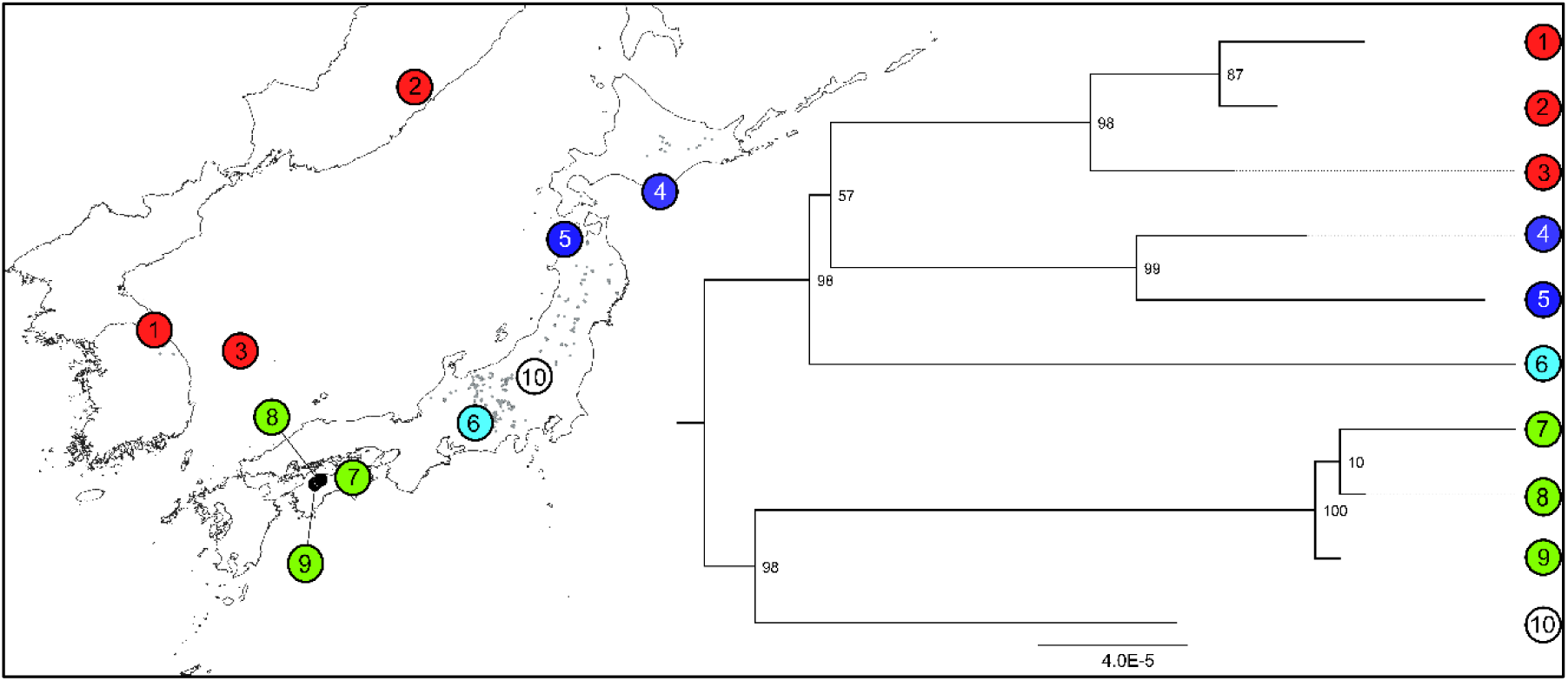
Phylogenetic tree based on the whole chloroplast genomes of *Rhododendron brachycarpum* assembled from samples from Japan, South Korea and Russia. The location of the identified clades are mapped. These results are based on the reference mapping to *R. shanii* (MW374796).

**Figure 12.**
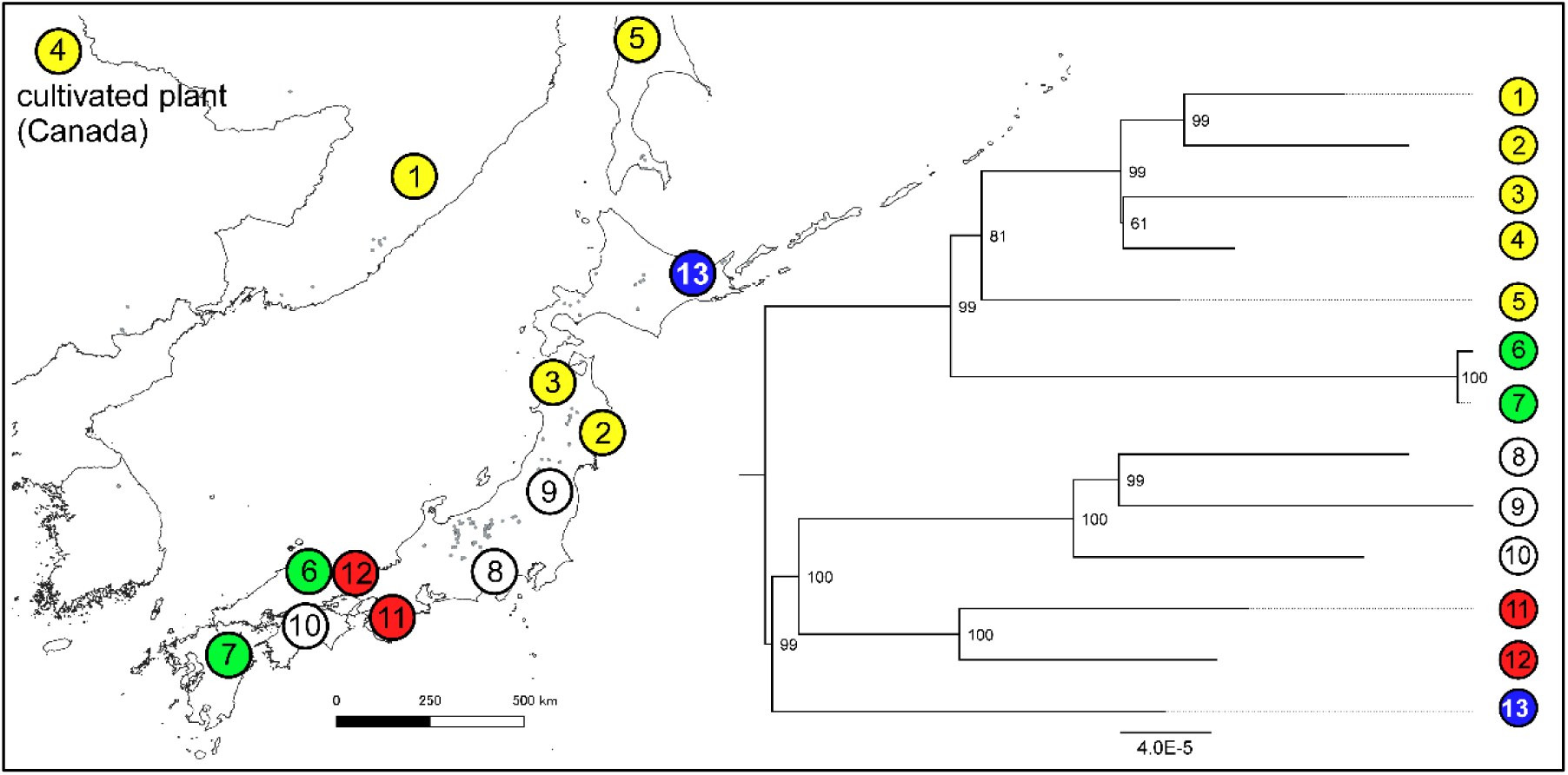
Phylogenetic tree based on the whole chloroplast genomes of *Vaccinium vitis-idaea* assembled from samples from Japan and Russia. The location of the identified clades are mapped. These results are based on reference mapping to the whole chloroplast genome of a cultivated individual of *Vaccinium vitis-idaea* from Canada.

## Discussion

This is one of the first studies to develop ultra barcodes for species representing a distinct biome and contributes to an increasing trend to assemble whole chloroplast genomes for genetic resource development studies. For example, Song *et al*. (2023) included whole chloroplast genomes, along with traditional short Sanger-based fragments, in a barcode library of the flowering plants of arid NW China while Krawczyk *et al*. (2018) used whole chloroplast genomes in a barcode library for the genus *Stipa*. However, studies investigating *within* species variation of the whole chloroplast genome remain rare. In this study the assembly of whole chloroplast genomes from short read genome skimming data facilitated the discovery of significant levels of intraspecific chloroplast variation in the 12 subalpine forest trees and shrubs. With the exception of *T. standishii*, this includes tens of SNPs, indels and polymorphic simple sequence repeat regions per species. A total of 31% and 41% of SNPs were parsimony informative for Japan and all samples, respectively. These will be useful for elucidating genetic relationships and divergence of populations across the range of these 12 species. However, hotspots of variation were exceedingly rare with the vast majority of variable genes or intragenic regions only having a single SNP.

While the whole chloroplast genomes of 10 species were efficiently and accurately assembled using *de novo* assembly-based methods this method failed using short reads for the two Ericaceae species, *R. brachycarpum* and *V*. *vitis-idaea*. However, we show that mapping reads to a reference even in the absence of a con-specific reference in the case of *R. brachycarpum* can provide reliable assay of chloroplast variation. For both species tens of SNPs and geographically based chloroplast lineages were identified.

### Potential applications in phylogeography and conservation genetics

By identifying chloroplast SNPs, indels and SSRs this study will accelerate phylogeographic and conservation genetic studies of these 12 important subalpine forests species. The whole chloroplast genome approach to identifying intraspecific chloroplast variation has distinct advantages over using methods based on previously published universal primers. Firstly, it maximises the ability to identify intraspecific chloroplast variation, whether it consists of rare singleton SNPs to deeply diverged lineages (Worth *et al*., 2021; Wang *et al*., 2023), in a genome that, as this study demonstrates, is mostly invariable at the species level (sites with SNPs comprised between 0.0015-0.107% of all sites in the 10 species where whole chloroplast genomes were assembled via *de novo* methods for Japan samples). In fact, any variation, especially potentially phylogenetically informative sites, were found to be scattered widely apart across the genome and therefore are not guaranteed to be found via traditional Sanger based methods. Secondly, primer pairs can be designed to target fragments that ensure the most efficient screening of chloroplast variation (both singletons and parsimony informative ones) according to each project’s resources and objectives. Thirdly, simultaneously, chloroplast SSRs which have a faster rate of evolution than other types of chloroplast polymorphism (Provan *et al*., 1999) and are particularly sensitive markers for assessing population size changes and genetic diversity (Provan, Powell and Hollingsworth, 2001) can be easily identified. Lastly, the method can reveal unexpected patterns of chloroplast sharing with congeneric species as demonstrated by the nesting of Genbank accessions of related species of *Betula* and *Berberis* in the intraspecific variation found in Japan of *B. ermanii* and *B. amurensis*.

The identification of areas with unique genetic lineages (i.e. evolutionary significant units (Moritz, 1994)) and/or high levels of genetic diversity could help to prioritise allocation of limited conservation resources and inform management decisions for the 12 subalpine species. This chloroplast information could also be used to identify seed source zones for reforestation or translocation (Tsumura, 2022), which is particularly important given the overall threat of decline of subalpine forests under global warming, deer browsing and the small and isolated nature of some populations. For example, *Ilex rugosa* has only one population on the island of Kyushu and one in the whole of the Chugoku area of western Japan, while the declining isolated population of *Oplopanax japonicus* on the Kii Peninsula depends almost entirely on the protection of deer proof fences for its persistence. However, in all these cases nothing is known about the divergence and genetic diversity of these populations. Range-wide studies are required to clarify the distribution of chloroplast variation identified in this study, including the significance of apparently southern versus northerly distributed distinct lineages identified in some of the 12 species.

### Genetic markers for ancient DNA studies

Fossils of subalpine plants including most of the 12 species investigated in this study are found in Last Glacial age sediments where they are sometimes abundant (Nishiuchi *et al*., 2017). Genetic studies that utilize the chloroplast genomes assembled in this study could contribute to studies of ancient DNA by providing an independent assessment of species diversity and abundance, e.g. via methods such as sedaDNA (Liu *et al*., 2020), to contrast with solely fossil based conclusions. Indeed, incorporating genetics into palaeoecological studies of Japan is an important endeavour because it could enable the identification of morphologically similar but ecologically diverged species from fossils. In Japan, conifers of the genera *Tsuga*, *Abies* and *Pinus* have both temperate and subalpine representatives while *Picea* has both geographically restricted and widespread representatives. Therefore, the inability to distinguish species of these genera when only fossil pollen is available and/or when macrofossils lack informative parts such as reproductive structures, has large implications for our understanding of past vegetation and migration/ range contraction histories of specific forest biomes. For conifers, whole chloroplast genomes are particularly promising for ancient DNA studies given the high level of species divergence of the chloroplast which is paternally inherited in conifers (Mogensen, 2009). In addition, having the whole chloroplast genome available means that any chloroplast genome fragment obtained from fossils or sediment using Next Generation Sequencing methods, typically comprising small fragments under 50 bp (Parducci *et al*., 2019), will likely be matchable to some part of the genome and, depending on the length and diversity of the fragment (and the level of chloroplast sharing in the case of angiosperms), the species will likely be identified. The likelihood of accurate species identification will only improve with increasing number of different chloroplast fragments recovered. In Japan, increasing the number of whole chloroplast genomes available especially for species rich genera (e.g. *Picea*) is crucial to increasing the potential and accuracy of ancient DNA studies. Knowledge about the intra-specific chloroplast lineages in each species may also provide an opportunity to investigate the past distribution of specific lineages.

## Supporting information

Vaccinium vitis-ideae reference mapping assembly

Rhododendron brachycarpum mapped_to_NC061396

Rhododendron brachycarpum mapped_to_MW374796

## Acknowledgements

We would like to thank C. Furusawa, H. Kanahara, Y. Kawamata and N. Takashima for assistance with laboratory work, A. Yorioka and T. Udino-Ihara for their invaluable assistance with obtaining sampling permission and all relevant national and prefectural agencies who provided sampling permission. In addition, we are grateful to K. Gyokusen, T. Katsuki, S. Kawano, Y. Nishiuchi, K. Obase, I. Tamaki, I. Tsuyama and K. Uchiyama for help in the field. This research was funded by the Japanese Society for the Promotion of Science KAKENHI grants 16H06197, 19H02980 and a Forestry and Forest Products Research Institute internal grant [number 201430] and the Korean Basic Science Institute (National Research Facilities and Equipment Center) of the Ministry of Education (grant number 2023R1A6C101B022).

## Data accessibility statement

All whole chloroplast genomes are available on Genbank for the ten species whose whole chloroplast genome was assembled using *de novo* methods while for *R. brachycarpum* and *V. vitis-idaea* results see the fasta alignments in the Supplementary Materials.

## Author contributions

The research was designed and performed by J.R.P.Worth, S. Kikuchi, S. Kanetani and S. Ueno. D. Takahashi, M. Aizawa, E. A. Marchuk, M. A. Polezhaeva, V. V. Sheiko and H. Jae Choi provided resources for this research. J.R.P.Worth and S. Ueno undertook the analyses and J.R.P.Worth interpreted the data and wrote the manuscript with input from all authors.

