## Supplementary Materials for "Chloroplast genome-based genetic resources for Japan’s threatened subalpine forests via genome skimming"

**Table S1** Information on each sample used for genome scanning and assembly of the whole chloroplast genome.

| No. | Species | DNA No. | Location | Latitude | Longitude | DNA source |
|---|---|---|---|---|---|---|
| 1 | *Abies veitchii* | Avi_46 | Mt Azuma, Fukushima, Japan | 37.70 | 140.25 | this study |
| 2 | *Abies veitchii* | Avi_76 | Lake Kirikomi, Tochigi, Japan | 36.82 | 139.44 | this study |
| 3 | *Abies veitchii* | Avi_116 | Mt Ena, Nagano, Japan | 35.44 | 137.61 | this study |
| 4 | *Abies veitchii* | Avi_163 | Mt Shakagatake, Nara, Japan | 34.12 | 135.91 | this study |
| 5 | *Abies veitchii* | Avi_98 | Mt Tsurugi, Tokushima, Japan | 33.85 | 134.11 | this study |
| 6 | *Abies veitchii* | Avi_31 | Mt Sasagamine, Ehime, Japan | 33.83 | 133.28 | this study |
| 7 | *Abies veitchii* | Avi_10 | Mt Ishizuchi, Ehime, Japan | 33.77 | 133.11 | this study |
| 8 | *Acer ukurunduense* | Acuk_137 | Lake Kussharo, Hokkaido, Japan | 43.63 | 144.27 | this study |
| 9 | *Acer ukurunduense* | Acuk_122 | Mt Iwaki, Aomori, Japan | 40.66 | 140.29 | this study |
| 10 | *Acer ukurunduense* | Acuk_183 | Mt Seorak, Gangwon, South Korea | 38.11 | 128.46 | this study |
| 11 | *Acer ukurunduense* | Acuk_16 | Yatsugatake Mtns, Nagano, Japan | 36.06 | 138.37 | this study |
| 12 | *Acer ukurunduense* | Acuk_39 | Mt Ena, Nagano, Japan | 35.44 | 137.61 | this study |
| 13 | *Acer ukurunduense* | Acuk_152 | Mt Misen, Nara, Japan | 34.18 | 135.91 | this study |
| 14 | *Acer ukurunduense* | Acuk_150 | Mt Misen, Nara, Japan | 34.18 | 135.91 | this study |
| 15 | *Berberis amurensis* | Ber_84 | Horoman Rv, Hokkaido, Japan | 42.11 | 143.05 | this study |
| 16 | *Berberis amurensis* | Ber_117 | Mt Byoubu, Miyagi, Japan | 38.11 | 140.45 | this study |
| 17 | *Berberis amurensis* | Ber_225 | Namyang, Ulleung Island, South Korea | 37.47 | 130.84 | this study |
| 18 | *Berberis amurensis* | Ber_33 | Yatsugatake Mtns, Nagano, Japan | 35.98 | 138.32 | this study |
| 19 | *Berberis amurensis* | Ber_69 | Mt Ouso, Okayama, Japan | 35.09 | 133.55 | this study |
| 20 | *Berberis amurensis* | Ber_50 | Mt Neko, Hiroshima, Japan | 35.04 | 133.19 | this study |
| 21 | *Berberis amurensis* | Ber_14 | Mt Higashiakaishi, Ehime, Japan | 33.87 | 133.37 | this study |
| 22 | *Berberis amurensis* | Ber_156 | Mt Shiraiwa, Miyazaki, Japan | 32.57 | 131.11 | this study |
| 23 | *Betula ermanii* | Bem_415 | Mt Krasnaya, Sakhalin Island, Russia | 46.93 | 142.84 | this study |
| 24 | *Betula ermanii* | Bem_218 | Mt Orofure, Hokkaido, Japan | 42.56 | 141.09 | this study |
| 25 | *Betula ermanii* | Bem_439 | Mt Iwaki, Aomori, Japan | 40.65 | 140.30 | this study |
| 26 | *Betula ermanii* | Bem_55 | Mt Azuma, Fukushima, Japan | 37.70 | 140.23 | this study |
| 27 | *Betula ermanii* | Bem_126 | Mt Hakusan, Ishikawa, Japan | 36.13 | 136.74 | this study |
| 28 | *Betula ermanii* | Bem_150 | Mt Ena, Nagano, Japan | 35.44 | 137.61 | this study |
| 29 | *Betula ermanii* | Bem_386 | Mt Jiri, Gyeongnam, South Korea | 35.32 | 127.70 | this study |
| 30 | *Betula ermanii* | Bem_544 | Mt Shakagatake, Nara, Japan | 34.12 | 135.91 | this study |
| 31 | *Betula ermanii* | Bem_89 | Mt Tsurugi, Tokushima, Japan | 33.85 | 134.11 | this study |
| 32 | *Betula ermanii* | Bem_15 | Mt Ishizuchi, Ehime, Japan | 33.75 | 133.16 | this study |
| 33 | *Ilex rugosa* | Iru_289 | Mt Mayorskaya, Sakhalin Island, Russia | 46.91 | 142.93 | this study |
| 34 | *Ilex rugosa* | Iru_255 | Yunipesotsu Rv., Hokkaido, Japan | 43.34 | 142.98 | this study |

| | | | | | | |
|---|---|---|---|---|---|---|
| 35 | *Ilex rugosa* | Iru_233 | Mt Iwaki, Aomori, Japan | 40.66 | 140.29 | this study |
| 36 | *Ilex rugosa* | Iru_Goyo | Mt Goyo, Iwate, Japan | 39.20 | 141.73 | this study |
| 37 | *Ilex rugosa* | Iru_110 | Mt Ena, Nagano, Japan | 35.45 | 137.62 | this study |
| 38 | *Ilex rugosa* | Iru_133 | Mt Asayama, Hiroshima, Japan | 34.78 | 132.37 | this study |
| 39 | *Ilex rugosa* | Iru_238 | Mt Odaigahara, Nara, Japan | 34.19 | 136.12 | this study |
| 40 | *Ilex rugosa* | Iru_247 | Mt Hakkyou, Nara, Japan | 34.17 | 135.91 | this study |
| 41 | *Ilex rugosa* | Iru_70 | Mt Higashiakaishi, Ehime, Japan | 33.88 | 133.37 | this study |
| 42 | *Ilex rugosa* | Iru_279 | Mt Ryoushi, Oita, Japan | 33.09 | 131.19 | this study |
| 43 | *Oplopanax japonicus* | Opj_137 | Mt Orofure, Hokkaido, Japan | 42.55 | 141.08 | this study |
| 44 | *Oplopanax japonicus* | Opj_150 | Mt Iwaki, Aomori, Japan | 40.66 | 140.28 | this study |
| 45 | *Oplopanax japonicus* | Opj_20 | Mt Azuma, Fukushima, Japan | 37.71 | 140.23 | this study |
| 46 | *Oplopanax japonicus* | Opj_60 | Mt Ena, Nagano, Japan | 35.44 | 137.61 | this study |
| 47 | *Oplopanax japonicus* | Opj_197 | Mt Hakkyou, Nara, Japan | 34.17 | 135.91 | this study |
| 48 | *Oplopanax japonicus* | Opj_204 | Mt Shakagatake, Nara, Japan | 34.12 | 135.91 | this study |
| 49 | *Oplopanax japonicus* | Opj_6 | Mt Ishizuchi, Ehime, Japan | 33.77 | 133.11 | this study |
| 50 | *Oplopanax japonicus* | Opj_15 | Mt Ishizuchi, Ehime, Japan | 33.77 | 133.11 | this study |
| 51 | *Picea jezoensis subsp. jezoensis* | KBR-183 | Khabarovsk, Khabarovsk, Russia | 48.30 | 135.10 | a |
| 52 | *Picea jezoensis subsp. jezoensis* | STK-11 | Shiretoko, Hokkaido, Japan | 43.92 | 145.07 | a |
| 53 | *Picea jezoensis subsp. hondoensis* | NIK-5 | Nikko, Tochigi, Japan | 36.75 | 139.44 | a |
| 54 | *Picea jezoensis subsp. hondoensis* | FUJ-3 | Mt Fuji, Yamanashi, Japan | 35.39 | 138.69 | a |
| 55 | *Picea jezoensis subsp. hondoensis* | ODI-13 | Mt Odaigahara, Nara, Japan | 34.18 | 136.11 | a |
| 56 | *Pinus koraiensis* | KUR 10-12 M | Kur-Urmi, Khabarovsk, Russia | 49.18 | 133.48 | b |
| 57 | *Pinus koraiensis* | MIN- SMS1 | Minami-Aizu, Tochigi, Japan | 36.98 | 139.55 | b |
| 58 | *Pinus koraiensis* | PAL 2-20 M | Mt Palgong, Gyeongsang, South Korea | 36.00 | 128.70 | b |
| 59 | *Pinus koraiensis* | NIS NSI22 | Mt Nishi-dake, Nagano, Japan | 35.95 | 138.34 | b |
| 60 | *Pinus koraiensis* | ONT OTK5 | Mt Ontake, Nagano, Japan | 35.92 | 137.53 | b |
| 61 | *Pinus koraiensis* | HIG HAK1 | Mt Higashiakaishi, Ehime, Japan | 33.87 | 133.38 | b |
| 62 | *Rhododendron brachycarpum* | Rbr_53 | Mt Ninomori, Ehime, Japan | 33.76 | 133.09 | this study |
| 63 | *Rhododendron brachycarpum* | Rbr_63 | Mt Ishizuchi, Ehime, Japan | 33.77 | 133.11 | this study |
| 64 | *Rhododendron brachycarpum* | Rbr_107 | Mt Kinsei, Nagano, Japan | 36.82 | 139.40 | this study |
| 65 | *Rhododendron brachycarpum* | Rbr_130 | Mt Tsurugi, Tokushima, Japan | 33.85 | 134.11 | this study |
| 66 | *Rhododendron brachycarpum* | Rbr_164 | Mt Ena, Nagano, Japan | 35.44 | 137.60 | this study |
| 67 | *Rhododendron brachycarpum* | Rbr_219 | Sikhote-Alin, Primorsky Krai, Russia | 45.13 | 135.87 | c |
| 68 | *Rhododendron brachycarpum* | Rbr_285 | Horoman Rv, Hokkaido, Japan | 42.10 | 143.04 | this study |
| 69 | *Rhododendron brachycarpum* | Rbr_317 | Mt Iwaki, Aomori, Japan | 40.66 | 140.29 | this study |
| 70 | *Rhododendron brachycarpum* | Rbr_391 | Hyeonpo, Ulleung Island, South Korea | 37.50 | 130.86 | c |
| 71 | *Rhododendron brachycarpum* | Rbr_403 | Mt Seorak, Gangwon, South Korea | 38.12 | 128.46 | this study |
| 72 | *Thuja standishii* | Kurobe_724 | Sakusawa Rv, Aomori, Japan | 40.51 | 140.35 | d |
| 73 | *Thuja standishii* | Kurobe_577 | Mt Sanpouiwa, Nagano, Japan | 36.26 | 136.84 | d |
| 74 | *Thuja standishii* | Kurobe_271 | Ooyamasawa, Saitama, Japan | 35.96 | 138.76 | d |
| 75 | *Thuja standishii* | Kurobe_438 | Mt Hanataka, Shimane, Japan | 35.41 | 132.75 | d |
| 76 | *Thuja standishii* | Kurobe_17 | Mt Higashiakaishi, Ehime, Japan | 33.87 | 133.37 | d |
| 77 | *Tsuga diversifolia* | 139 | Mt Iwaki, Aomori, Japan | 40.66 | 140.29 | this study |
| 78 | *Tsuga diversifolia* | 1279 | Mt Azuma, Fukushima, Japan | 37.71 | 140.25 | e |
| 79 | *Tsuga diversifolia* | 1399 | Mt Shakagatake, Nara, Japan | 34.11 | 135.90 | this study |

| | | | | | | |
|---|---|---|---|---|---|---|
| 80 | *Tsuga diversifolia* | 1748 | Mt Ishizuchi, Ehime, Japan | 33.76 | 133.13 | this study |
| 81 | *Vaccinium vitis-idaea* | Vvi_Goyo | Mt Goyo, Iwate, Japan | 39.20 | 141.72 | this study |
| 82 | *Vaccinium vitis-idaea* | Vvi_29 | Mt Azuma, Fukushima, Japan | 37.70 | 140.24 | this study |
| 83 | *Vaccinium vitis-idaea* | Vvi_58 | Mt Higashiakaishi, Ehime, Japan | 33.88 | 133.38 | this study |
| 84 | *Vaccinium vitis-idaea* | Vvi_98 | Mt Fuji, Yamanashi, Japan | 35.39 | 138.71 | this study |
| 85 | *Vaccinium vitis-idaea* | Vvi_172 | Mt Mimata, Oita, Japan | 33.10 | 131.24 | this study |
| 86 | *Vaccinium vitis-idaea* | Vvi_185 | Mt Hyono, Hyogo, Japan | 35.35 | 134.51 | this study |
| 87 | *Vaccinium vitis-idaea* | Vvi_237 | Mt Iwaki, Aomori, Japan | 40.65 | 140.30 | this study |
| 88 | *Vaccinium vitis-idaea* | Vvi_258 | Bihoro Pass, Hokkaido, Japan | 43.65 | 144.25 | this study |
| 89 | *Vaccinium vitis-idaea* | Vvi_280 | Mt Hakkyou, Nara, Japan | 34.17 | 135.91 | this study |
| 90 | *Vaccinium vitis-idaea* | Vvi_323 | Taulanka Rv, Sakhalin Island, Russia | 50.43 | 142.65 | this study |
| 91 | *Vaccinium vitis-idaea* | Vvi_343 | Snezhny Stream, Primorsky Krai, Russia | 46.37 | 136.49 | this study |
| 92 | *Vaccinium vitis-idaea* | Vvi_347 | Kabutogasen, Tottori, Japan | 35.39 | 133.58 | this study |

**Table S2** Details of the raw Illumina HiSeq2000 data, whole chloroplast genome assembly and Genbank accession numbers for each sample.

| Species | Sample | Read count | Read Length (bp) | Size of cp genome | % of the chloroplast genome in the WGS data | Average base coverage | Chloroplast whole genome Genbank Accession No. |
|---|---|---|---|---|---|---|---|
| *Abies veitchii* | Avi_10 | 8172102 | 150 | 121412 | 0.70 | 71 | OR459817 |
| *Abies veitchii* | Avi_31 | 8225070 | 150 | 121319 | 0.63 | 64 | OR459818 |
| *Abies veitchii* | Avi_46 | 8186790 | 150 | 121352 | 1.06 | 107 | OR459819 |
| *Abies veitchii* | Avi_76 | 8209876 | 150 | 121391 | 0.35 | 36 | OR459820 |
| *Abies veitchii* | Avi_98 | 8233300 | 150 | 121382 | 0.40 | 41 | OR459821 |
| *Abies veitchii* | Avi_116 | 8256456 | 150 | 121298 | 0.24 | 24 | OR459822 |
| *Abies veitchii* | Avi_163 | 8247702 | 150 | 121403 | 0.73 | 74 | OR459823 |
| *Acer ukurunduense* | Acuk_16 | 8057932 | 150 | 156478 | 1.32 | 101.6 | OR178638 |
| *Acer ukurunduense* | Acuk_39 | 8055730 | 150 | 156526 | 2.72 | 210.1 | OR346893 |
| *Acer ukurunduense* | Acuk_122 | 8028980 | 150 | 156717 | 1.03 | 79 | OR346894 |
| *Acer ukurunduense* | Acuk_137 | 8066992 | 150 | 156634 | 1.41 | 109.1 | OR346895 |
| *Acer ukurunduense* | Acuk_150 | 7408814 | 150 | 156461 | 2.00 | 142 | OR346896 |
| *Acer ukurunduense* | Acuk_152 | 8054010 | 150 | 156486 | 1.70 | 130.9 | OR346897 |
| *Acer ukurunduense* | Acuk_183 | 8161446 | 150 | 156555 | 3.28 | 256.4 | OR346898 |
| *Berberis amurensis* | Ber_14 | 7560572 | 150 | 166564 | 3.28 | 223 | OR415855 |
| *Berberis amurensis* | Ber_33 | 8020712 | 150 | 166581 | 2.66 | 192.2 | OR415856 |
| *Berberis amurensis* | Ber_50 | 8155834 | 150 | 166699 | 1.95 | 143.4 | OR415857 |
| *Berberis amurensis* | Ber_69 | 8143652 | 150 | 166601 | 4.29 | 314.5 | OR415858 |
| *Berberis amurensis* | Ber_84 | 8146410 | 150 | 166277 | 0.92 | 67.3 | OR415859 |
| *Berberis amurensis* | Ber_117 | 8236476 | 150 | 166547 | 1.91 | 141.6 | OR415860 |
| *Berberis amurensis* | Ber_156 | 8042662 | 150 | 166708 | 1.11 | 80.4 | OR415861 |
| *Berberis amurensis* | Ber_225 | 8219058 | 150 | 166652 | 1.71 | 126.4 | OR415862 |
| *Betula ermanii* | Bem_15 | 8072038 | 150 | 160616 | 5.08 | 382.6 | OR346907 |
| *Betula ermanii* | Bem_55 | 8081786 | 150 | 160525 | 4.91 | 370.6 | OR346908 |
| *Betula ermanii* | Bem_89 | 8096638 | 150 | 160624 | 6.08 | 459.5 | OR346909 |
| *Betula ermanii* | Bem_126 | 8110514 | 150 | 160628 | 4.02 | 304.5 | OR346910 |
| *Betula ermanii* | Bem_150 | 8081738 | 150 | 160623 | 2.91 | 219.9 | OR346911 |
| *Betula ermanii* | Bem_218 | 8082198 | 150 | 160554 | 1.61 | 121.2 | OR350408 |
| *Betula ermanii* | Bem_386 | 8126662 | 150 | 160564 | 5.61 | 426 | OR350409 |
| *Betula ermanii* | Bem_415 | 8153942 | 150 | 160474 | 6.64 | 506.3 | OR350410 |
| *Betula ermanii* | Bem_439 | 8068662 | 150 | 160504 | 0.66 | 49.7 | OR350411 |
| *Betula ermanii* | Bem_544 | 8067292 | 150 | 160616 | 1.85 | 139.3 | OR350412 |
| *Ilex rugosa* | Iru_Mt_Goyo | 9376860 | 150 | 157583 | 5.73 | 511 | OR415849 |
| *Ilex rugosa* | Iru_70 | 8161498 | 150 | 157584 | 4.95 | 384.2 | OR350413 |
| *Ilex rugosa* | Iru_110 | 8036618 | 150 | 157588 | 6.02 | 460.2 | OR350414 |
| *Ilex rugosa* | Iru_133 | 8084876 | 150 | 157588 | 4.43 | 341 | OR350415 |
| *Ilex rugosa* | Iru_233 | 8208476 | 150 | 157586 | 5.05 | 394.5 | OR415854 |
| *Ilex rugosa* | Iru_238 | 8128704 | 150 | 157587 | 6.28 | 485.9 | OR415853 |

| | | | | | | | |
|---|---|---|---|---|---|---|---|
| *Ilex rugosa* | Iru_247 | 8203264 | 150 | 157586 | 5.27 | 411.3 | OR415852 |
| *Ilex rugosa* | Iru_255 | 8126582 | 150 | 157588 | 6.53 | 505.2 | OR415851 |
| *Ilex rugosa* | Iru_279 | 8182844 | 150 | 157584 | 5.07 | 394.8 | OR415850 |
| *Ilex rugosa* | Iru_289 | 8078470 | 150 | 157589 | 5.86 | 450.9 | OR634941 |
| *Oplopanax japonicus* | Opj_6 | 8233174 | 150 | 156135 | 1.51 | 119.1 | OR346899 |
| *Oplopanax japonicus* | Opj_15 | 8065620 | 150 | 156136 | 2.38 | 184.7 | OR346900 |
| *Oplopanax japonicus* | Opj_20 | 8033934 | 150 | 156171 | 5.19 | 400.1 | OR346901 |
| *Oplopanax japonicus* | Opj_60 | 8028276 | 150 | 156147 | 0.74 | 56.8 | OR346902 |
| *Oplopanax japonicus* | Opj_137 | 8051298 | 150 | 156173 | 2.24 | 173.4 | OR346903 |
| *Oplopanax japonicus* | Opj_150 | 8304698 | 150 | 156157 | 3.20 | 255.3 | OR346904 |
| *Oplopanax japonicus* | Opj_197 | 8065004 | 150 | 156129 | 5.44 | 421.9 | OR346905 |
| *Oplopanax japonicus* | Opj_204 | 8150510 | 150 | 156109 | 2.43 | 190.6 | OR346906 |
| *Picea jezoensis* subsp. *jezoensis* | KBR-183 | 8291156 | 150 | 124146 | 1.54 | 154.4 | OR528865 |
| *Picea jezoensis* subsp. *jezoensis* | STK-11 | 8230048 | 150 | 124152 | 0.88 | 87.9 | OR528868 |
| *Picea jezoensis* subsp. *hondoensis* | NIK-5 | 8298352 | 150 | 124307 | 0.72 | 71.9 | OR528866 |
| *Picea jezoensis* subsp. *hondoensis* | FUJ-3 | 8302864 | 150 | 124254 | 1.36 | 136 | OR522708 |
| *Picea jezoensis* subsp. *hondoensis* | ODI-13 | 8275008 | 150 | 124395 | 0.98 | 97.7 | OR528867 |
| *Pinus koraiensis* | KUR 10-12 M | 10142646 | 150 | 116924 | 2.44 | 316.9 | OR506266 |
| *Pinus koraiensis* | MIN- SMS1 | 8276368 | 150 | 117010 | 0.54 | 57.7 | OR506270 |
| *Pinus koraiensis* | PAL 2-20 M | 8332358 | 150 | 116995 | 1.91 | 204 | OR501833 |
| *Pinus koraiensis* | NIS NSI22 | 7145684 | 150 | 117103 | 0.54 | 49 | OR506268 |
| *Pinus koraiensis* | ONT OTK5 | 7184380 | 150 | 116955 | 0.71 | 65.2 | OR506269 |
| *Pinus koraiensis* | HIG HAK1 | 7251034 | 150 | 116986 | 0.52 | 47.9 | OR506267 |
| *Thuja standishii* | Kurobe_17 | 8228298 | 150 | 130925 | 1.16 | 109.8 | OR583020 |
| *Thuja standishii* | Kurobe_271 | 8590264 | 150 | 130772 | 0.88 | 86.6 | OR574833 |
| *Thuja standishii* | Kurobe_438 | 8086554 | 150 | 130638 | 0.75 | 69.4 | OR583021 |
| *Thuja standishii* | Kurobe_577 | 8393284 | 150 | 130855 | 1.18 | 113.2 | OR583022 |
| *Thuja standishii* | Kurobe_724 | 8312736 | 150 | 130661 | 0.88 | 84.3 | OR583023 |
| *Tsuga diversifolia* | 1748 | 6780430 | 100 | 120925 | 0.85 | 47.5 | OR528869 |
| *Tsuga diversifolia* | 139 | 6750072 | 100 | 121029 | 0.83 | 46.1 | OR522709 |
| *Tsuga diversifolia* | 1399 | 6776732 | 100 | 120963 | 0.66 | 36.9 | OR528870 |
| *Tsuga diversifolia* | 1279 | 26472610 | 100 | 121043 | 0.94 | 206.5 | MH171102 |
| *Rhododendron brachycarpum* | Rbr_53 | 8008090 | 150 | - | - | - | - |
| *Rhododendron brachycarpum* | Rbr_63 | 8020002 | 150 | - | - | - | - |
| *Rhododendron brachycarpum* | Rbr_107 | 8027746 | 150 | - | - | - | - |
| *Rhododendron brachycarpum* | Rbr_130 | 8051922 | 150 | - | - | - | - |
| *Rhododendron brachycarpum* | Rbr_164 | 7689294 | 150 | - | - | - | - |
| *Rhododendron brachycarpum* | Rbr_219 | 8224694 | 150 | - | - | - | - |
| *Rhododendron brachycarpum* | Rbr_285 | 8054172 | 150 | - | - | - | - |
| *Rhododendron brachycarpum* | Rbr_317 | 8118370 | 150 | - | - | - | - |
| *Rhododendron brachycarpum* | Rbr_391 | 8115156 | 150 | - | - | - | - |
| *Rhododendron brachycarpum* | Rbr_403 | 8011840 | 150 | - | - | - | - |
| *Vaccinium vitis-idaea* | Vvi_MtGoyo | 8233942 | 150 | - | - | - | - |
| *Vaccinium vitis-idaea* | Vvi_29 | 8115424 | 150 | - | - | - | - |
| *Vaccinium vitis-idaea* | Vvi_58 | 8070266 | 150 | - | - | - | - |

| | | | | | | | |
|---|---|---|---|---|---|---|---|
| *Vaccinium vitis-idaea* | Vvi_98 | 8189094 | 150 | - | - | - | - |
| *Vaccinium vitis-idaea* | Vvi_172 | 8089282 | 150 | - | - | - | - |
| *Vaccinium vitis-idaea* | Vvi_185 | 8124396 | 150 | - | - | - | - |
| *Vaccinium vitis-idaea* | Vvi_237 | 8066678 | 150 | - | - | - | - |
| *Vaccinium vitis-idaea* | Vvi_258 | 8122258 | 150 | - | - | - | - |
| *Vaccinium vitis-idaea* | Vvi_280 | 8213386 | 150 | - | - | - | - |
| *Vaccinium vitis-idaea* | Vvi_323 | 8051532 | 150 | - | - | - | - |
| *Vaccinium vitis-idaea* | Vvi_343 | 8013730 | 150 | - | - | - | - |
| *Vaccinium vitis-idaea* | Vvi_347 | 8208628 | 150 | - | - | - | - |

**Table S3** Summary of genetic diversity identified in the *de novo* assembled whole chloroplast genomes for the six non-Japanese endemic study species including single nucleotide polymorphisms and indels based on the all samples-dataset. Excluding *Ilex rugosa*, *Rhododendron brachycarpum* and *Vaccinium vitis-idaea* the samples from outside Japan include Genbank accessions.

| Species | No. samples | Total sites | Sites (excluding gaps) | Monomorphic sites | Singletons | Parsimony informative sites | Nucleotide diversity (*Pi*) | Average number of nucleotide differences (*k*) | Indel events | InDel Diversity per site (*Pi*(i)) |
|---|---|---|---|---|---|---|---|---|---|---|
| *Berberis amurensis* | 12 | 129855 | 128938 | 128789 | 90 | 59 | 0.0003 | 39.0 | 156 | 0.0004 |
| *Betula ermanii* | 11 | 135262 | 133763 | 133662 | 75 | 26 | 0.00019 | 25.7 | 208 | 0.00046 |
| *Ilex rugosa* | 10 | 131510 | 131475 | 131446 | 21 | 8 | 0.00006 | 8.1 | 30 | 0.00009 |
| *Acer ukurunduense* | 8 | 130441 | 129432 | 129368 | 12 | 52 | 0.0002 | 25.6 | 105 | 0.00033 |
| *Picea jezoensis* | 6 | 124676 | 123842 | 123640 | 131 | 71 | 0.00068 | 84.6 | 92 | 0.00031 |
| *Pinus koraiensis* | 7 | 117229 | 117225 | 116257 | 23 | 17 | 0.00014 | 16.0 | 65 | 0.00022 |
| *Rhododendron brachycarpum*[1] | 10 | 157211 | - | 157150 | 53 | 54 | 0.00022 | - | - | - |
| *Rhododendron brachycarpum*[2] | 10 | 155954 | - | 155830 | 142 | 44 | 0.00019 | - | - | - |
| *Vaccinium vitis-idaea*[3] | 13 | 144322 | - | 144084 | 154 | 84 | 0.00039 | - | - | - |

[1] mapped to MW374796; [2] mapped to OM373082; [3] mapped to the *V. vitis-idaea* whole chloroplast genome assembled from the Oxford Nanopore and Illumina NovaSeq reads Hirabayashi, Debnath and Owens (2023)

**Table S4** Summary of genetic diversity identified at simple sequence repeat regions in the *de novo* assembled whole chloroplast genomes of the six non-Japanese endemic species based on the all samples-dataset. Excluding *Ilex rugosa*, the samples from outside Japan include Genbank accessions.

| Species | No. samples | No. polymorphic SSRs (mono-repeats >9 length) | No. monomorphic SSRs (mono-repeats >9 length) | No. polymorphic SSRs (di-repeats >9 length) | No. monomorphic SSRs (di-repeats >9 length) | No. polymorphic SSRs (tri-repeats >9 length) | No. monomorphic SSRs (tri-repeats >9 length) |
|---|---|---|---|---|---|---|---|
| *Berberis amurensis* | 12 | 55 | 17 | 1 | 9 | 2 | 2 |
| *Betula ermanii* | 11 | 48 | 15 | 4 | 13 | 2 | 11 |
| *Ilex rugosa* | 10 | 21 | 30 | 0 | 4 | 0 | 5 |
| *Acer ukurunduense* | 8 | 45 | 30 | 0 | 7 | 2 | 21 |
| *Picea jezoensis* | 6 | 26 | 1 | 4 | 6 | 0 | 4 |
| *Pinus koraiensis* | 7 | 28 | 11 | 0 | 6 | 0 | 0 |

**Table S5** The top ten most variable regions of the *de novo* assembled whole chloroplast genome for six non-Japanese endemic species according to nucleotide diversity (*Pi*) including samples from within and outside Japan. Excluding *Ilex rugosa*, the samples from outside Japan include Genbank accessions. Intergenic spacers are italicized while genes are shown in bold.

| Rank | *Acer ukurunduense* | *Berberis amurensis* | *Betula ermanii* | *Ilex rugosa* | *Picea jezoensis* | *Pinus koraiensis* |
|---|---|---|---|---|---|---|
| 1 | *rps19—rpl2* | *psbI—trnS-GCU* | *psbZ—trnG-GCC* | *ndhG—ndhI* | *rpl2—rpl23* | *rps12—rps7* |
| 2 | *infA—rps8* | *trnH-GUG—psbA* | *rps15—ycf1* | *psbI—trnS-GCU* | *rpoA—rps11* | *trnT-UGU—rps4* |
| 3 | *ndhD—psaC* | **accD** | *ndhH—rps15* | *trnH-GUG—psbA* | *clpP_rps12* | *trnfM-CAU—trnG-GCC* |
| 4 | *rpl16—rps3* | *ycf1—ndhF* | **rps15** | *rps4—trnT-UGU* | **trnR—UCU** | **psbK** |
| 5 | *trnH-GUG—psbA* | *ndhF—rpl32* | *ndhD—psaC* | *trnS-UGA—psbZ* | **trnS—UGA** | *rrn5—trnR-ACG* |
| 6 | *rpl33—rps18* | *rbcL—accD* | **rpl14** | *trnG-GCC—trnfM-CAU* | *trnW-CCA—petG* | *rpl23—trnI-GAU* |
| 7 | *trnW-CCA—trnP-UGG* | *trnL-UAG—ccsA* | *trnN-GUU— ycf1* | *psbB—psbT* | *ycf12—clpP* | *trnQ-UUG—psbK* |
| 8 | *rps8—rpl14* | *atpI—rps2* | *petG—trnW-CCA* | *rps2—rpoC2* | *chlB—trnQ-UUG* | *trnS-GCU—psaM* |
| 9 | *ndhC—trnV-UAC* | *trnQ-UUG—psbK* | *trnD-GUC—trnY-GUA* | *psbA—matK* | *trnH-GUG—trnT-GGU* | *psbK—psbI* |
| 10 | *ycf4—cemA* | *trnS-GGA—rps4* | *ndhF—rpl32* | *rpoB_trnC—trnC-GCA* | *ycf12—psbB* | **ndhK** |

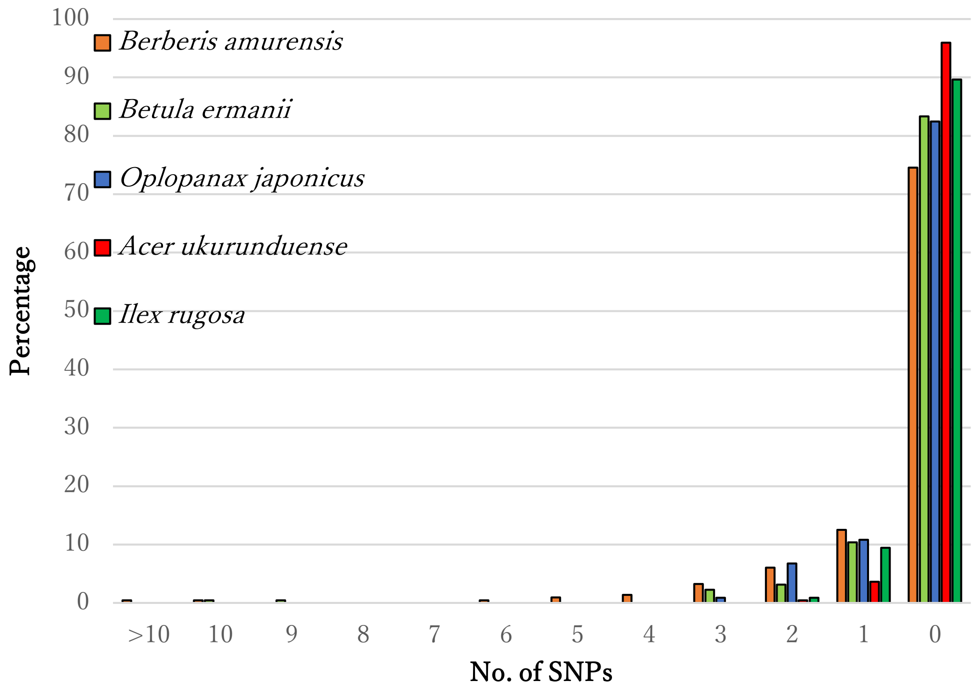


**Figure S1** The percentage of chloroplast regions (genes and intergenic spacers) without SNPs and with SNPs (from 1 to over 10 per region) for the 5 angiosperm species where whole chloroplast genomes could be assembled by the *de novo* method. These results are based on the only Japan dataset.

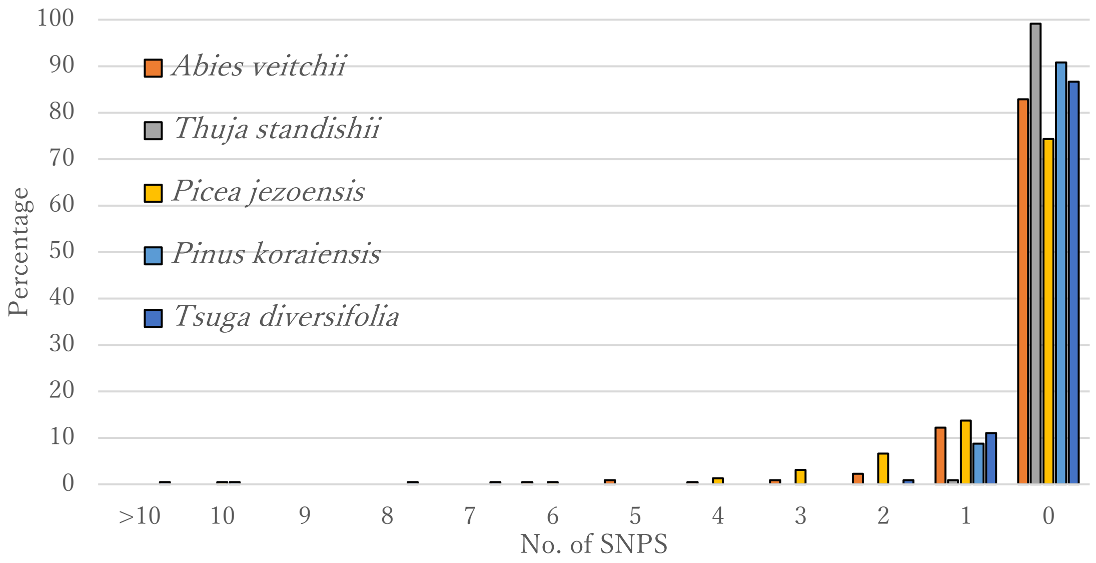


**Figure S2** The percentage of chloroplast regions (genes and intergenic spacers) without SNPs and with SNPs (from 1 to over 10 per region) for the 5 conifers species where whole chloroplast genomes could be assembled by the *de novo* method. These results are based on the only Japan dataset.

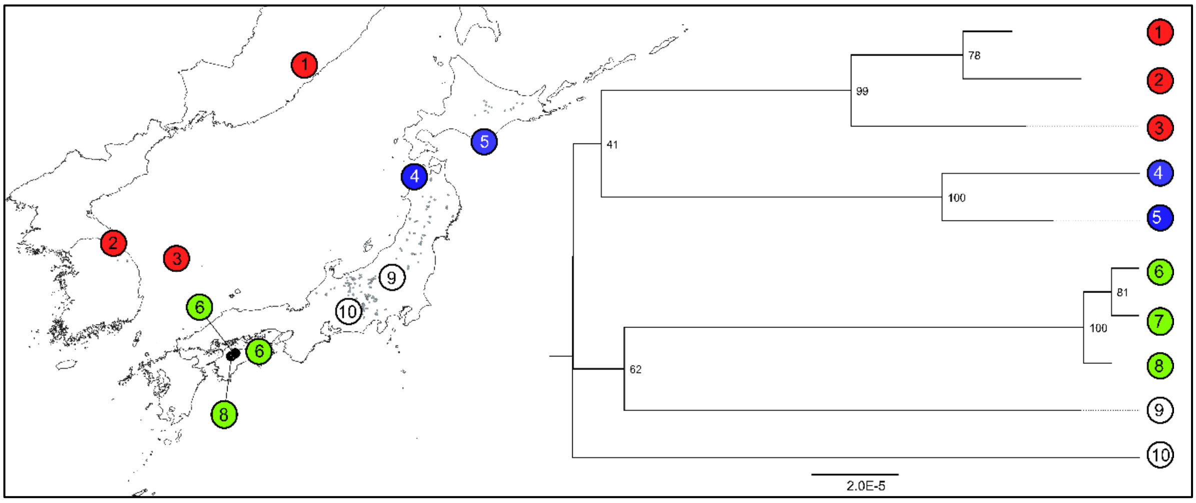


**Figure S3** Phylogenetic tree based on the whole chloroplast genomes of *Rhododendron brachycarpum* assembled from sampled from Japan, South Korea and Russia. The location of the identified clades are mapped. These are the results based on the reference mapping to *R. calophytum* (OM373082).
